# A comparison shopper’s guide to forest datasets

**DOI:** 10.1101/2025.06.02.657445

**Authors:** Lucy G. Lee, Valerie J. Pasquarella, Benjamin Glass, Luca L. Morreale, Nina Chung, Xiaojie Gao, Jonathan R. Thompson

## Abstract

Recent advances in remote sensing, data availability, and cloud-based computing have led to a rapid expansion of publicly accessible datasets characterizing forest cover and land use. These datasets are widely used in ecological research, natural resource management, and policymaking. However, the sheer number of available products—and the often-subtle differences among them—pose significant challenges for users seeking the most appropriate dataset for their specific objectives. Here, we evaluate 27 derived products from 12 publicly available sources that quantify tree or forest cover and use across the conterminous United States (CONUS), with temporal coverage ranging from 5 to 30 years. The products include both satellite-based remote sensing data and ground-based national forest inventory data. We ask: How, why, and where do these datasets differ in their estimates of forest extent and change over time? Our analysis reveals that estimates of total forest area at the CONUS scale differ by over 2,000,000 km², and correlations among forest change estimates vary widely in both direction and statistical significance. To support dataset selection and interpretation, we developed an open-access map comparison tool using Google Earth Engine. Our findings highlight the substantial implications of dataset choice for understanding forest dynamics and underscore the need for careful selection and transparent reporting in forest-related analyses.

## Introduction

Forests are a critical part of the Earth system. They sustain human lives in myriad ways, from mitigating climate change, to conserving biodiversity, to provisioning clean air, water, food, fuel, and fiber (FAO, 2024). To ensure that forests and the benefits they provide are protected, policy makers and resource managers need reliable estimates of how and where forest cover is changing (Keenan et al., 2015). Estimates of changes in forest extent directly inform policies regarding forest carbon stocks and fluxes (e.g., Harris et al., 2021; Pötzschner et al., 2022) and programs designed to mitigate climate change through avoided deforestation (Teo et al., 2023). Managing future timber supplies similarly requires reliable estimates of changes in forest extent through time (Bousfield et al., 2023). International programs aimed at reducing forest loss and degradation rely on national forest area estimates, and these estimates affect the funding for which countries are eligible (Chazdon et al., 2016). Forest maps are essential for estimating the location and magnitude of forest fragmentation (Morreale et al., 2024), with concomitant effects on ecosystem health and functioning (Watson et al., 2018), and are often the only tool available to assess the impact of policies aimed at reducing undesirable forest changes over large scales (e.g., Ramirez-Reyes et al., 2018). The primary data challenge faced by those in need of accurate information on forest area and change is not one of data scarcity; rather, modern consumers of forest area information are often faced with an abundance of forest maps and datasets to choose from, each with their own assumptions and idiosyncrasies. Here we compare derivative products from twelve different forest and tree datasets and attempt to help end-users, including data analysts and policy-makers, understand their relative strengths and weaknesses and emphasize key points of differentiation that can help with dataset selection.

Historically, estimates of forest extent were derived from field sampling and/or interpretation of aerial photographs (Moessner, 1953). Today, most estimates of forest area are derived from satellite remote sensing, which has been used since the early 1970s to map land cover and land-cover change (e.g., Fassnacht et al., 2024; Gómez et al., 2016; Hemati et al., 2021; Loveland, 2012). A primary advantage of using satellite imagery for forest monitoring is its availability over large areas with regular revisit cycles on the order of days to weeks, thus providing large amounts of data over space and time. Comparisons of remote sensing-based forest maps show that spatial resolution (Wickham & Riitters, 2019), class definitions (Congalton et al., 2014; Estoque et al., 2018), and classification errors (Congalton et al., 2014; Dong et al., 2012; Estoque et al., 2018) all contribute to differences in resulting maps. Such methodological characteristics are important considerations in any project utilizing forest maps, in addition to the fundamental conceptual question that must underpin any study of forests and forest change – what constitutes “forest”?

“Forest” as a descriptive label can refer to a location’s land use or land cover (Coulston et al., 2014; Woodall et al., 2016). Land *cover* describes directly observable landscape characteristics, i.e., forest cover is the area of land that exceeds a defined threshold of tree canopy cover (Nedd et al., 2021). In contrast, land *use* describes what humans use the land for, i.e., its management intent. While an area classified as forest or tree land cover is frequently also classified as forest land use, this is not always the case. For example, a recently clear-cut forest would not be classified as forest cover (instead it may be classified as barren, grass, or shrub) while this same area would be classified as forest land use, because its management intent is to remain forested as it regrows. Additionally, neither forest cover nor use encompass all tree cover. Trees outside forests – such as those in urban or agricultural areas – may provide the same benefits as trees in forests (Arroyo-Rodríguez et al., 2020; Global Forest Review, 2023; Liu et al., 2023) but are often classified as the broader land cover or use in which they live (e.g., developed, residential, agriculture). Whether forest is interpreted as a land use or a land cover (or something else) can have significant consequences, as it forms the foundation which guides policy and management (Chazdon et al., 2016). Multiple studies (e.g., Chazdon et al., 2016; Coulston et al., 2014; Woodall et al., 2016) have asserted that the different uses and values of forests require different definitions, including distinctions based on land cover and land use. Nonetheless, land use and cover are frequently conflated (e.g., Estoque et al., 2022; Watson et al., 2018; Winkler et al., 2021), likely owing to the different methods needed to assess each and the difficulty of integrating consistent land-use data into classifications over large, heterogeneous areas (Tulbure et al., 2022).

Today, forest cover (i.e., forest land cover) is typically mapped across a grid of pixels using satellite remote sensing based on the presence of trees (and sometimes other vegetation) within each pixel, often with threshold requirements of percent coverage or canopy height (Table 1). Consistency of definitions across datasets and associated classification errors are consistently identified as two primary sources of disagreement among forest-cover maps (Chen et al., 2020; Congalton et al., 2014; Estoque et al., 2018). Because forest land cover is defined based on the presence of trees, forest-cover maps are sensitive to disturbance-driven changes in land cover, such as tree harvest or wildfire. Furthermore, the spatial resolution (pixel size) of the remote sensing data determines the minimum size that a cluster of trees must be to be classified as forest, which directly affects assessments of forest connectivity and fragmentation (Hernando et al., 2017; Morreale et al., 2024; Wickham & Riitters, 2019). Land cover definitions of forest also typically do not distinguish between monoculture plantations and native forests (Chazdon et al., 2016; Van Holt et al., 2016), although they are compositionally and ecologically distinct (Hall et al., 2012; C. Wang et al., 2022), which may limit utility for analysis requiring this distinction.

**Table 1.** Definitions of forest or tree cover by dataset.

| Dataset | Forest/Tree Class Definitions | Related publications |
| --- | --- | --- |
| Dynamic World | **Trees:** “Any significant clustering of dense vegetation, typically with a closed or dense canopy. Taller and darker than surrounding vegetation (if surrounded by other vegetation). For example, wooded vegetation, dense green shrubs, cluster of dense, tall vegetation within savannas, plantations such as apples, bananas, citrus, and rubber, swamp (dense/tall vegetation with no obvious water), or any mix or burned areas of these.” | (Brown et al., 2022) |
| US EPA Greenhouse Gas Inventory | **Forest Land:** “A land-use category that includes areas at least 120 feet (36.6 meters) wide and at least one acre (0.4 hectare) in size with at least 10 percent cover (or equivalent stocking) by live trees including land that formerly had such tree cover and that will be naturally or artificially regenerated. Trees are woody plants having a more or less erect perennial stem(s) capable of achieving at least 3 inches (7.6 cm) in diameter at breast height, or 5 inches (12.7 cm) diameter at root collar, and a height of 16.4 feet (5 m) at maturity in situ. Forest Land includes all areas recently having such conditions and currently regenerating or capable of attaining such condition in the near future. Forest Land also includes transition zones, such as areas between forest and non-forest lands that have at least 10 percent cover (or equivalent stocking) with live trees and forest areas adjacent to urban and built-up lands. Unimproved roads and trails, streams, and clearings in forest areas are classified as forest if they are less than 120 feet (36.6 m) wide or an acre (0.4 ha) in size. However, land is not classified as Forest Land if completely surrounded by urban or developed lands, even if the criteria are consistent with the tree area and cover requirements for Forest Land. These areas are classified as Settlements. In addition, Forest Land does not include land that is predominantly under an agricultural land use (Nelson et al., 2020).” | (Nelson et al., 2020; U.S. Environmental Protection Agency, 2023; Walters et al., 2023) |
| ESA WorldCover | **Tree Cover:** “This class includes any geographic area dominated by trees with a cover of 10% or more. Other land cover classes (shrubs and/or herbs in the understorey, built-up, permanent water bodies, ...) can be present below the canopy, even with a density higher than trees. Areas planted with trees for afforestation purposes and plantations (e.g. oil palm, olive trees) are included in this class. This class also includes tree covered areas seasonally or permanently flooded with fresh water except for mangroves.” | (Van De Kerchove et al., 2020) |
| ESRI Global Land Use Land Cover | **Trees:** “Any significant clustering of tall (~15-m or higher) dense vegetation, typically with a closed or dense canopy; examples: wooded vegetation, clusters of dense tall vegetation within savannas, plantations, swamp or mangroves (dense/tall vegetation with ephemeral water or canopy too thick to detect water underneath).” | (Karra et al., 2021) |
| Forest Inventory and Analysis (FIA) | **Forest Land:** “Forest land has at least 10 percent canopy cover of trees of any size, or has had at least 10-percent canopy cover of trees in the past, based on the presence of stumps, snags, or other evidence, and that will be naturally or artificially regenerated. Additionally, the land is not subject to nonforest use(s) that prevent normal tree regeneration and succession, such as regular mowing, intensive grazing, or recreation | (USDA Forest Service, 2023) |
|  | activities. Forest land includes transition zones, such as areas between heavily forested and nonforested lands that are at least 10-percent canopy cover with trees and forest areas adjacent to urban and built-up lands. Also included are pinyon-juniper and chaparral areas in the West and afforested areas. The minimum area for classification of forest land is 1 acre (0.4 ha) in size and 120 feet (36.6 m) wide measured stem-to-stem from the outer-most edge. Unimproved roads and trails, streams, and clearings in forest areas are classified as forest if less than 120 feet wide. This is the domestic reporting definition which is different from the definition used in international reporting (which includes a minimum tree canopy height criteria).”<br>**Timberland:** “Unreserved forest land capable of producing 20 cubic feet per acre per year of wood from trees classified as timber species (see “Timber species”) and designated as a timber forest type.” |  |
| Hansen Global Forest Cover | **Treecover2000:** “Tree canopy cover for year 2000, defined as canopy closure for all vegetation taller than 5m in height.”<br>**Lossyear:** “Year of gross forest cover loss event. Forest loss during the study period, defined as a stand-replacement disturbance, or a change from a forest to non-forest state. Encoded as either 0 (no loss) or else a value in the range 1–20, representing loss detected primarily in the year 2001–2020, respectively.” | (Hansen et al., 2013) |
| Land Change Monitoring, Assessment, and Projection (LCMAP) | **Tree Cover:** “Tree-covered land where the tree cover density is greater than 10%. Cleared or harvested trees (i.e., clearcuts) will be mapped according to current cover (e.g., Barren, Grass/Shrub).”<br>**Wetland:** “Lands where water saturation is the determining factor in soil characteristics, vegetation types, and animal communities. Wetlands are composed of mosaics of water, bare soil, and herbaceous or wooded vegetated cover.” | (US Geological Survey, 2022) |
| Land Change Monitoring System (LCMS) Land Cover | “**1. TREES:** Live or standing dead trees. **3. SHRUBS:** Shrubs. **4. GRASS/FORB/HERBACEOUS:** Perennial grasses, forbs, or other forms of herbaceous vegetation. **5. BARREN OR IMPERVIOUS:** 1) Bare soil exposed by disturbance (e.g., soil uncovered by mechanical clearing or forest harvest), as well as perennially barren areas such as deserts, playas, rock outcroppings (including minerals and other geologic materials exposed by surface mining activities), sand dunes, salt flats, and beaches. Roads made of dirt and gravel are also considered barren or 2) man-made materials that water cannot penetrate, such as paved roads, rooftops, and parking lots.” | (Housman, Heyer, et al., 2023) |
| LCMS Land Use | “**FOREST:** Land that is planted or naturally vegetated and which contains (or is likely to contain) 10% or greater tree cover at some time during a near-term successional sequence. This may include deciduous, evergreen and/or mixed categories of natural forest, forest plantations, and woody wetlands.” | (Housman, Heyer, et al., 2023) |
| MODIS Land Cover<br>Type 1:<br>International Geosphere- | **Land Cover Type 1 and Type 2 class descriptions**<br>“**1. Evergreen Needleleaf Forests:** dominated by evergreen conifer trees (canopy >2m). Tree cover >60%. **2. Evergreen Broadleaf Forests:** dominated by evergreen broadleaf and palmate trees (canopy >2m). Tree cover >60%. **3. Deciduous Needleleaf Forests:** dominated by deciduous needleleaf (larch) trees (canopy >2m). Tree cover >60%. **4. Deciduous** | (Sulla-Menashe & Friedl, 2022) |
| Biosphere Programme (IGBP) classification<br>Type 2: University of Maryland (UMD) classification<br>Type 3: Leaf Area Index (LAI) classification<br>Type 4: BIOME-Biogeochemical Cycles (BGC) classification<br>Type 5: Plant Functional Types (PFT) classification | **Broadleaf Forests:** dominated by deciduous broadleaf trees (canopy >2m). Tree cover >60%. **5. Mixed Forests:** dominated by neither deciduous nor evergreen (40-60% of each) tree type (canopy >2m). Tree cover >60%.”<br>*Land Cover Type 3 class descriptions*<br>“**5. Evergreen Broadleaf Forests:** dominated by evergreen broadleaf and palmate trees (canopy >2m). Tree cover >60%. **6. Deciduous Broadleaf Forests:** dominated by deciduous broadleaf trees (canopy >2m). Tree cover >60%. **7. Evergreen Needleleaf Forests:** dominated by evergreen conifer trees (canopy >2m). Tree cover >60%. **8. Deciduous Needleleaf Forests:** dominated by deciduous needleleaf (larch) trees (canopy >2m). Tree cover >60%.”<br>*Land Cover Type 4 class descriptions*<br>“**1. Evergreen Needleleaf Vegetation:** dominated by evergreen conifer trees and shrubs (>1m). Woody vegetation cover >10%. **2. Evergreen Broadleaf Vegetation:** dominated by evergreen broadleaf and palmate trees and shrubs (>1m). Woody vegetation cover >10%. **3. Deciduous Needleleaf Vegetation:** dominated by deciduous needleleaf (larch) trees and shrubs (>1m). Woody vegetation cover >10%. **4. Deciduous Broadleaf Vegetation:** dominated by deciduous broadleaf trees and shrubs (>1m). Woody vegetation cover >10%.”<br>*Land Cover Type 5 class descriptions*<br>“**1. Evergreen Needleleaf Trees:** dominated by evergreen conifer trees (>2m). Tree cover >10%. **2. Evergreen Broadleaf Trees:** dominated by evergreen broadleaf and palmate trees (>2m). Tree cover >10%. **3. Deciduous Needleleaf Trees:** dominated by deciduous needleleaf (larch) trees (>2m). Tree cover >10%. **4. Deciduous Broadleaf Trees:** dominated by deciduous broadleaf trees (>2m). Tree cover >10%.” |  |
| National Land Cover Database (NLCD) Land Cover | “**Deciduous forest:** areas dominated by trees generally greater than 5 meters tall, and greater than 20% of total vegetation cover. More than 75% of the tree species shed foliage simultaneously in response to seasonal change.<br>**Evergreen forest:** areas dominated by trees generally greater than 5 meters tall, and greater than 20% of total vegetation cover. More than 75% of the tree species maintain their leaves all year. Canopy is never without green foliage.<br>**Mixed forest:** areas dominated by trees generally greater than 5 meters tall, and greater than 20% of total vegetation cover. Neither deciduous nor evergreen species are greater than 75% of total tree cover.<br>**Woody wetlands:** areas where forest or shrubland vegetation accounts for greater than 20% of vegetative cover and the soil or substrate is periodically saturated with or covered with water.” | (Jin et al., 2019; U.S. Geological Survey, 2024) |
| NLCD Tree Canopy Cover | “Percent of the pixel that's covered by tree canopy. No masking of obvious non-tree areas is performed for this product.” | (Housman, Bender, et al., 2023) |
| National Resources Inventory (NRI) | “**Forest Land:** A Land cover/use category that is at least 10 percent stocked by single-stemmed woody species of any size that will be at least 4 meters (13 feet) tall at maturity. Also included is land bearing evidence of natural regeneration of tree cover (cut over forest or abandoned | (USDA Natural Resources Conservation Service, 2015) |
|  | farmland) and not currently developed for non-forest use. Ten percent stocked, when viewed from a vertical direction, equates to an areal canopy cover of leaves and branches of 25 percent or greater. The minimum area for classification as forest land is 1 acre, and the area must be at least 100 feet wide.” |  |

Forest as a land use can be similarly based on the presence of a certain density and/or size of trees, but it also can include areas without trees if they will be regenerated (Chazdon et al., 2016; Table 3). One of the challenges with mapping forest land use is understanding management intent, as “forests” from a use perspective cannot always be directly observed and management intent is sometimes unknown or unknowable (Woodall et al., 2016). Like forest cover, definitions of forest land use frequently include a minimum area requirement (see Table 1), which constrains the minimum size of a “forest”. However, forest-cover and forest-use datasets have different relationships to space and time. Forest-use data aims to assess intended use within a longer, often multi-year, time frame compared to forest cover, which assesses current conditions at a snapshot in time (Coulston et al., 2014; Winkler et al., 2021). Forest-use data is often more limited in spatial detail compared to forest-cover data (Coulston et al., 2014; Winkler et al., 2021) and is typically available less frequently (i.e., coarser temporal resolution) because it involves more labor intensive data collection methods, often including field-based measurements. Forest-use data, including national forest inventories, collected through field measurements often include other variables such as tree species and size, and it may be possible to distinguish tree plantations from natural forests. While most traditional forest-use datasets rely on field measurements, advances of remote sensing data in recent decades has made remote sensing-based forest-use data available, such as the USDA’s Landscape Change Monitoring System (LCMS). Such data utilizes time-series data and algorithms to identify “fluctuating state” changes that estimate forest use from land-cover change cycles, and distinguish these areas from persistent loss of tree cover (see Pasquarella et al., 2022).

Despite sharing a common ontology, use- and cover-based forest datasets frequently disagree on whether forests are expanding or declining. In the eastern US, studies have found that forest use and cover are not strongly correlated (Coulston et al., 2014; Woodall et al., 2016). Oloffson and others (2016) found continuous deforestation in the New England states based on Landsat time series data (i.e., land cover) but their results did not align in either direction or magnitude of change with trends documented in the widely-used U.S. Forest Service Forest Inventory Analysis (FIA), a land-use dataset. Regional studies in the U.S. have documented loss of forest cover (Drummond & Loveland, 2010; Olofsson et al., 2016; Woodall et al., 2016), increases in forest cover and use (Coulston et al., 2014), and forest-cover trends that change over time, with periods of loss and gain in the same study period (Adams et al., 2019). National research has documented increases in forest use (Nelson et al., 2020; Sleeter et al., 2018) and decreases in forest cover at a national scale (Homer et al., 2020; Nelson et al., 2020; Sleeter et al., 2018). Moreover, an international analysis by Holmgren (2015) found that countries containing half the world’s forests – including Russia, China, India, Canada, Australia, and the United States – show strong disagreements in forest use versus cover. Most commonly, forest use is found to increase or remain stable while forest cover decreases, due in large part to forestry practices which alter land cover but not land use (Curtis et al., 2018; Holmgren, 2015; Woodall et al., 2016), as well as farmland abandonment in naturally forested areas (Sleeter et al., 2018). Conflicting results inevitably lead to uncertainty about whether current forest management and other land-use policies are effective and what policies should be implemented in the future.

Which dataset is best depends on the question the data is intended to answer, and users need to understand the assumptions and characteristics behind datasets in order to match the dataset to their question, as well as statistical methods required to correct biases in map-based estimates. While other studies have considered small numbers of forest or land cover datasets (e.g., Coulston et al., 2014, Woodall et al. 2016, Congalton et al., 2014, Estoque et al., 2018, Chen et al., 2020) or the connection between forest definitions and datasets (e.g., Chazdon et al., 2016, Zalles et al., 2014), here we provide the first analysis of a large suite of open access forest datasets with the goal of aiding dataset selection. In this study, we compile and compare existing forest datasets within the conterminous United States (CONUS) to quantify their variation and provide a foundation for discussion of their characteristics and differences in an applied context. We examine 27 products derived from twelve different datasets (nine remote sensing-based datasets and three inventory-based datasets; Table 2) to answer the following questions:

1. How, why, and where do commonly used datasets vary in their estimates of forest area?
2. How, why, and where do commonly used datasets vary in their estimates of forest area change over time?
3. What tools and guidance can be offered to users of forest maps to help them identify appropriate datasets for specific applications?

**Table 2.**
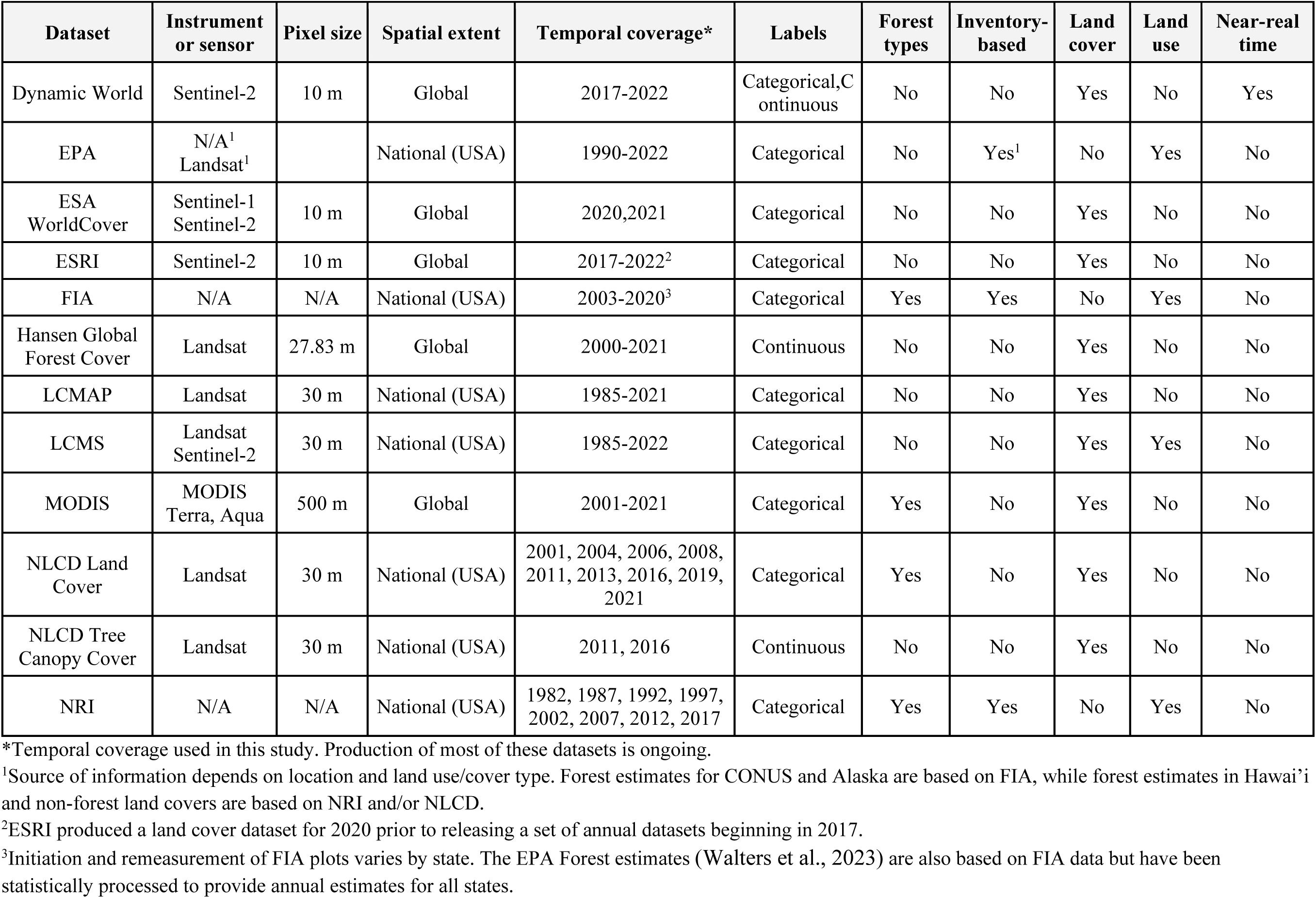
Characteristics of the forest and tree cover datasets analyzed in this study.

## Methods

We focus on open access datasets that are readily available on the Google Earth Engine platform, either in the main Earth Engine Data Catalog (https://developers.google.com/earth-engine/datasets) or via the Earth Engine Community Catalog (Roy et al., 2023; https://gee-community-catalog.org/). Table 3 records the classes used in each dataset. The dataset configurations described in Table 3 are better understood with Table 1, which describes each dataset’s definition(s) of forest use/cover or tree cover.

**Table 3.**
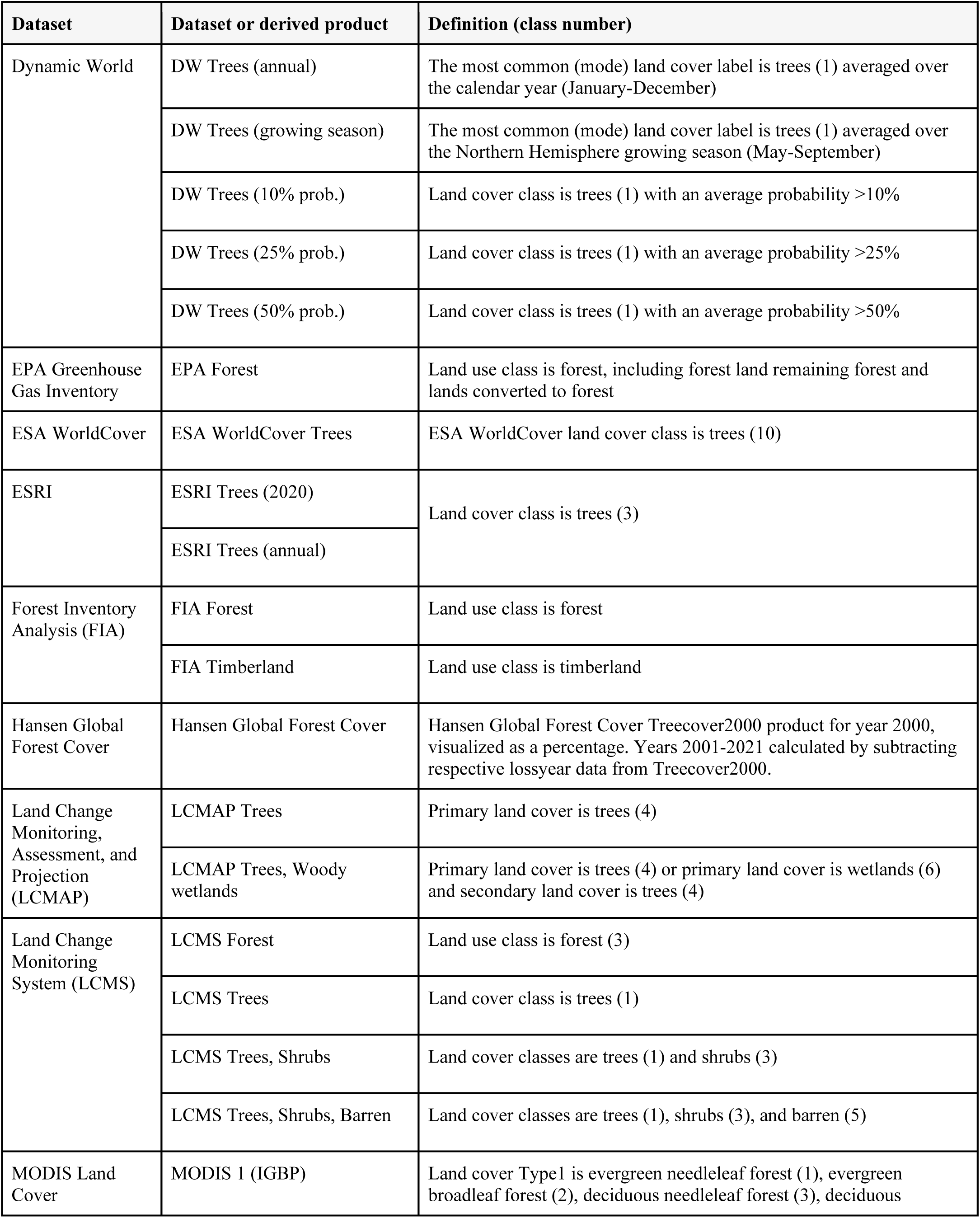
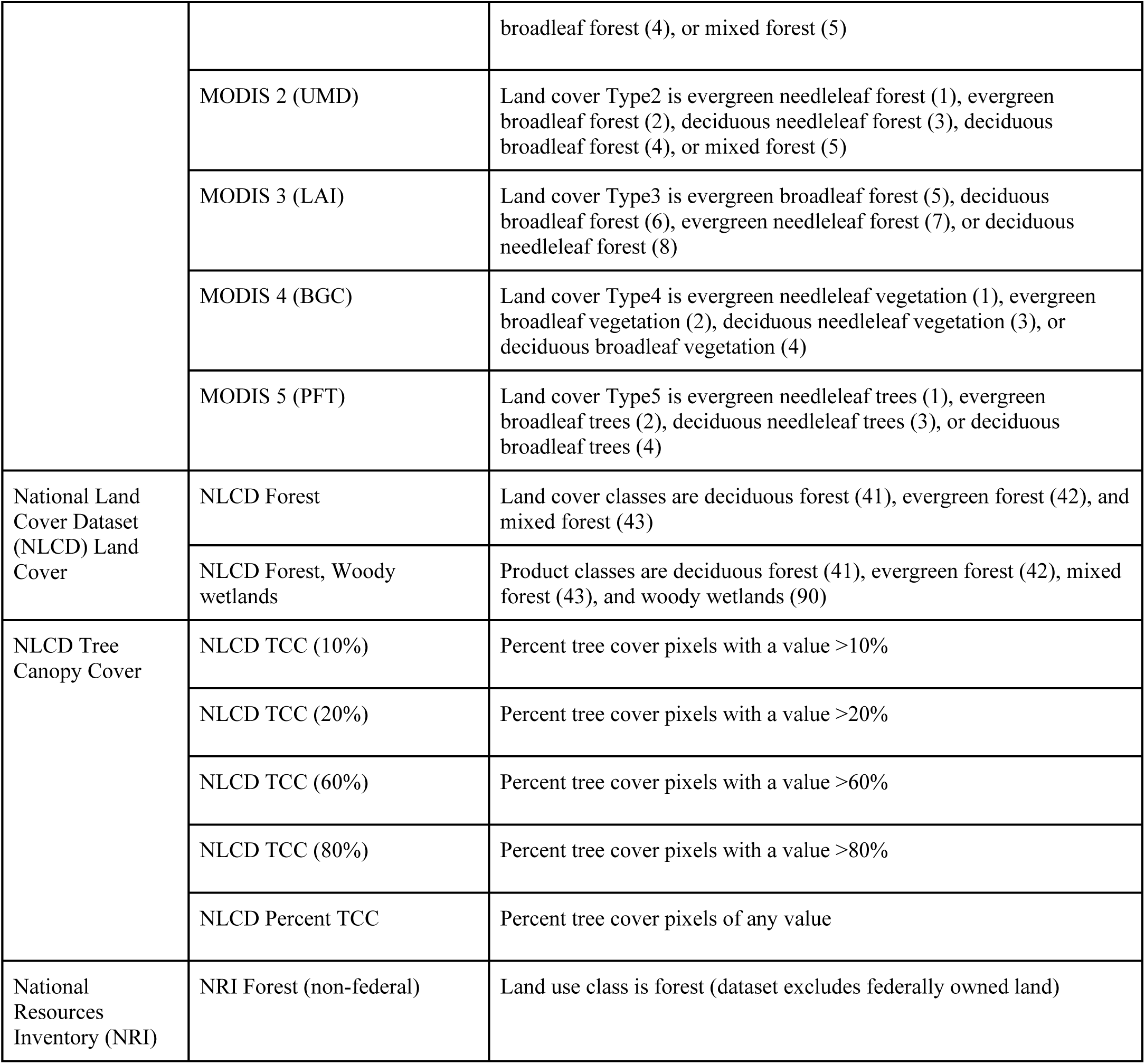
Summary of dataset configurations analyzed.

| Dataset | Dataset or derived product | Definition (class number) |
| --- | --- | --- |
| Dynamic World | DW Trees (annual) | The most common (mode) land cover label is trees (1) averaged over the calendar year (January-December) |
|  | DW Trees (growing season) | The most common (mode) land cover label is trees (1) averaged over the Northern Hemisphere growing season (May-September) |
|  | DW Trees (10% prob.) | Land cover class is trees (1) with an average probability >10% |
|  | DW Trees (25% prob.) | Land cover class is trees (1) with an average probability >25% |
|  | DW Trees (50% prob.) | Land cover class is trees (1) with an average probability >50% |
| EPA Greenhouse Gas Inventory | EPA Forest | Land use class is forest, including forest land remaining forest and lands converted to forest |
| ESA WorldCover | ESA WorldCover Trees | ESA WorldCover land cover class is trees (10) |
| ESRI | ESRI Trees (2020) | Land cover class is trees (3) |
|  | ESRI Trees (annual) |  |
| Forest Inventory Analysis (FIA) | FIA Forest | Land use class is forest |
|  | FIA Timberland | Land use class is timberland |
| Hansen Global Forest Cover | Hansen Global Forest Cover | Hansen Global Forest Cover Treecover2000 product for year 2000, visualized as a percentage. Years 2001-2021 calculated by subtracting respective lossyear data from Treecover2000. |
| Land Change Monitoring, Assessment, and Projection (LCMAP) | LCMAP Trees | Primary land cover is trees (4) |
|  | LCMAP Trees, Woody wetlands | Primary land cover is trees (4) or primary land cover is wetlands (6) and secondary land cover is trees (4) |
| Land Change Monitoring System (LCMS) | LCMS Forest | Land use class is forest (3) |
|  | LCMS Trees | Land cover class is trees (1) |
|  | LCMS Trees, Shrubs | Land cover classes are trees (1) and shrubs (3) |
|  | LCMS Trees, Shrubs, Barren | Land cover classes are trees (1), shrubs (3), and barren (5) |
| MODIS Land Cover | MODIS 1 (IGBP) | Land cover Type1 is evergreen needleleaf forest (1), evergreen broadleaf forest (2), deciduous needleleaf forest (3), deciduous |
|  |  | broadleaf forest (4), or mixed forest (5) |
|  | MODIS 2 (UMD) | Land cover Type2 is evergreen needleleaf forest (1), evergreen broadleaf forest (2), deciduous needleleaf forest (3), deciduous broadleaf forest (4), or mixed forest (5) |
|  | MODIS 3 (LAI) | Land cover Type3 is evergreen broadleaf forest (5), deciduous broadleaf forest (6), evergreen needleleaf forest (7), or deciduous needleleaf forest (8) |
|  | MODIS 4 (BGC) | Land cover Type4 is evergreen needleleaf vegetation (1), evergreen broadleaf vegetation (2), deciduous needleleaf vegetation (3), or deciduous broadleaf vegetation (4) |
|  | MODIS 5 (PFT) | Land cover Type5 is evergreen needleleaf trees (1), evergreen broadleaf trees (2), deciduous needleleaf trees (3), or deciduous broadleaf trees (4) |
| National Land Cover Dataset (NLCD) Land Cover | NLCD Forest | Land cover classes are deciduous forest (41), evergreen forest (42), and mixed forest (43) |
|  | NLCD Forest, Woody wetlands | Product classes are deciduous forest (41), evergreen forest (42), mixed forest (43), and woody wetlands (90) |
| NLCD Tree Canopy Cover | NLCD TCC (10%) | Percent tree cover pixels with a value >10% |
|  | NLCD TCC (20%) | Percent tree cover pixels with a value >20% |
|  | NLCD TCC (60%) | Percent tree cover pixels with a value >60% |
|  | NLCD TCC (80%) | Percent tree cover pixels with a value >80% |
|  | NLCD Percent TCC | Percent tree cover pixels of any value |
| National Resources Inventory (NRI) | NRI Forest (non-federal) | Land use class is forest (dataset excludes federally owned land) |

In addition to remote sensing datasets, we also include state-level summaries from three inventory-based estimates for the United States in our comparisons. These include the US Department of Agriculture (USDA) Forest Inventory and Analysis (FIA) forest census (Bechtold & Patterson, 2005), the US Environmental Protection Agency (EPA) estimates of forest area (Walters et al., 2023), and the USDA National Resources Inventory (NRI).

We use all data available, except for the 1992 National Land Cover Dataset (NLCD), which is not comparable with later years (Jin et al., 2019). FIA is unique among the datasets analyzed because it is the only dataset for which all states do not share the same years of available data. While we use all available FIA data, due to varying years of plot remeasurement across states, CONUS-level trend analysis is only possible from 2011 through 2019 (see Table S2).

For remote sensing-based datasets, we used Google Earth Engine to pre-process raster layers and calculate pixel-based estimates of total forest area by state. All datasets were first converted to binary masks where “non-forest”=0 and “forest”=1 using one or more categorical remappings to capture potential variability in class definitions. We then apply a reducer to get the total count of “forest” pixels per year within each state using the US Census 2018 TIGER States dataset to define state boundaries. We reproject all datasets to a common projection (US Albers, EPSG:5070) while retaining native resolution for consistency across pixel area calculations.

For FIA data, we used the R package *rFIA* (Stanke et al., 2020) to estimate the annual area of forested land and timberlands at the state-level. The *rFIA* package provides access to the latest inventory data and facilitates estimation of forest attributes following standard FIA protocols.

To account for variability in proportion of forested area by state, all forest area estimates were normalized by state land area from the 2010 US Census (US Census Bureau, 2010), except the NRI, which excludes federally owned land from forest area estimates. To calculate normalized NRI forest estimates, we used the Protected Areas Database of the United States (U.S. Geological Survey Gap Analysis Project, 2022) to calculate the area of lands owned in fee by the US federal government and subtracted this area from each state’s total land area.

Tabular outputs from remote sensing dataset processing and other tabular data sources datasets were combined and analyzed in R (version 4.2.1). Simple linear regressions and Pearson’s correlation coefficients were calculated using all available data between 2000-2019 to maximize temporal overlap across datasets. We included datasets that have at least four observations over this period. Some datasets do not have full coverage over this time period (see Table 2). Statistical tests were considered significant at 95% confidence.

Links to datasets used are in Table S1. Code for the Earth Engine portion of analysis (i.e., summarizing remote sensing datasets) is available at https://github.com/valpasq/ee-forests/tree/main. Code for the statistical tests and figures in this manuscript is available at https://github.com/hf-thompson-lab/forest-comparison.

## Results

### Large discrepancies in forest extent at the CONUS scale

Estimates of forest extent, as well as the direction, magnitude, significance, and agreement of trends in forest extent vary substantially among datasets at the CONUS and state scales. There is a 32.9% difference between the highest and lowest estimate of “forest” in 2020 (Fig. 1). Most datasets estimate the total tree- or forest-covered CONUS area between 2,000,000 km^2^ and 3,000,000 km^2^ (Fig. 1), a range that is more than twice the size of California. The EPA’s estimate, which is used for international reporting, lands in the middle, at approximately 2,500,000 km^2^ (Fig. 1). The EPA Forest estimates are primarily derived from plots measured in the field by the FIA program, and have a unique definition wherein forests that are completely surrounded by development are not classified as forest but rather as “settlements” (Table 1), possibly contributing to the lower EPA estimates compared to FIA for the years when these datasets overlap (Fig. 1).

**Figure 1.**
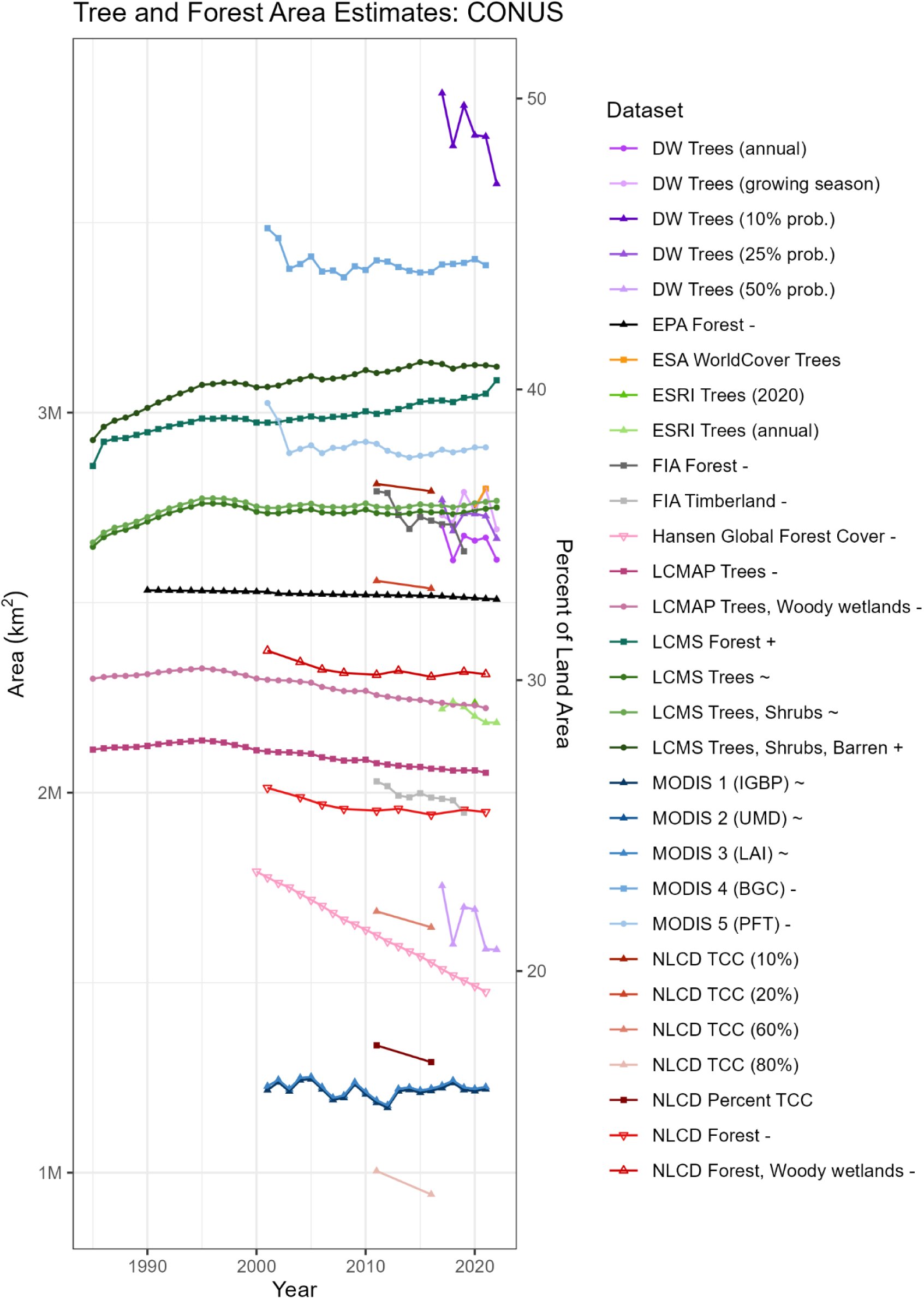
Forest or tree area over time, in square kilometers and as a percent of land area, for the conterminous United States. Symbols following dataset names, if present, indicate whether the dataset had a significantly increasing (+), significantly decreasing (-), or insignificant (∼) trend from 2000-2019. NRI is not included in this figure because it excludes federally owned forests (see Figures S1 and S2).

Type classifications provided by the MODIS Land-Cover Type (MCD12Q1) Version 6.1 data product differed by approximately 2,000,000 km^2^, more than four times the size of the state of California, due to algorithmic differences as well as differences in definitions of “forest” (Fig. 1; Table 1). MODIS 4 (BGC) definition of forest has the lowest canopy height criteria and includes shrubs in addition to trees (Table 1), which helps explain why this dataset has the largest CONUS forest estimates out of all datasets analyzed (Fig. 1). Conversely, MODIS 1-3 have the lowest CONUS forest estimates (Fig. 1). These datasets share the same canopy height criteria as MODIS 5 (PFT), but require >60% tree cover for classification as forest, the highest coverage threshold of all datasets analyzed (Table 1).

Forest estimates from LCMS are greater than NLCD estimates by more than 1,000,000 km^2^ (Fig. 1), even though both datasets are generated from 30-meter Landsat data. NLCD requires 20% tree cover for classification as forest, double the 10% coverage threshold used by most datasets analyzed here, including LCMS (Table 1). LCMS includes areas not currently meeting that threshold but that are “likely” to contain 10% tree cover in the future (i.e., it includes clearcuts and conceives of forest as a land use) whereas NLCD does not (Table 1). These definitional differences in both the threshold of canopy coverage and whether “forest” means a land use or a land cover may interact to produce such disparate estimates.

### Variation in interannual change estimates

Due to the large range in area estimates among datasets, interannual variability within each dataset is difficult to discern in Figure 1. When comparing change from previous measurement within each dataset, differences in the magnitude of change and interannual variability become apparent (Fig. 2). EPA Forest data stand out for its relatively consistent low magnitude change (but note that the loss of 4,700 km^2^ between 2001 and 2002 appears to be an outlier in this dataset; Fig. 2). In contrast, mode aggregations from Dynamic World have a high long-term magnitude of change and high interannual variability: the loss of ∼91,000 km^2^ from 2017 to 2018 is followed by a gain of ∼64,000 km^2^ from 2018 to 2019 (Fig. 2).

**Figure 2.**
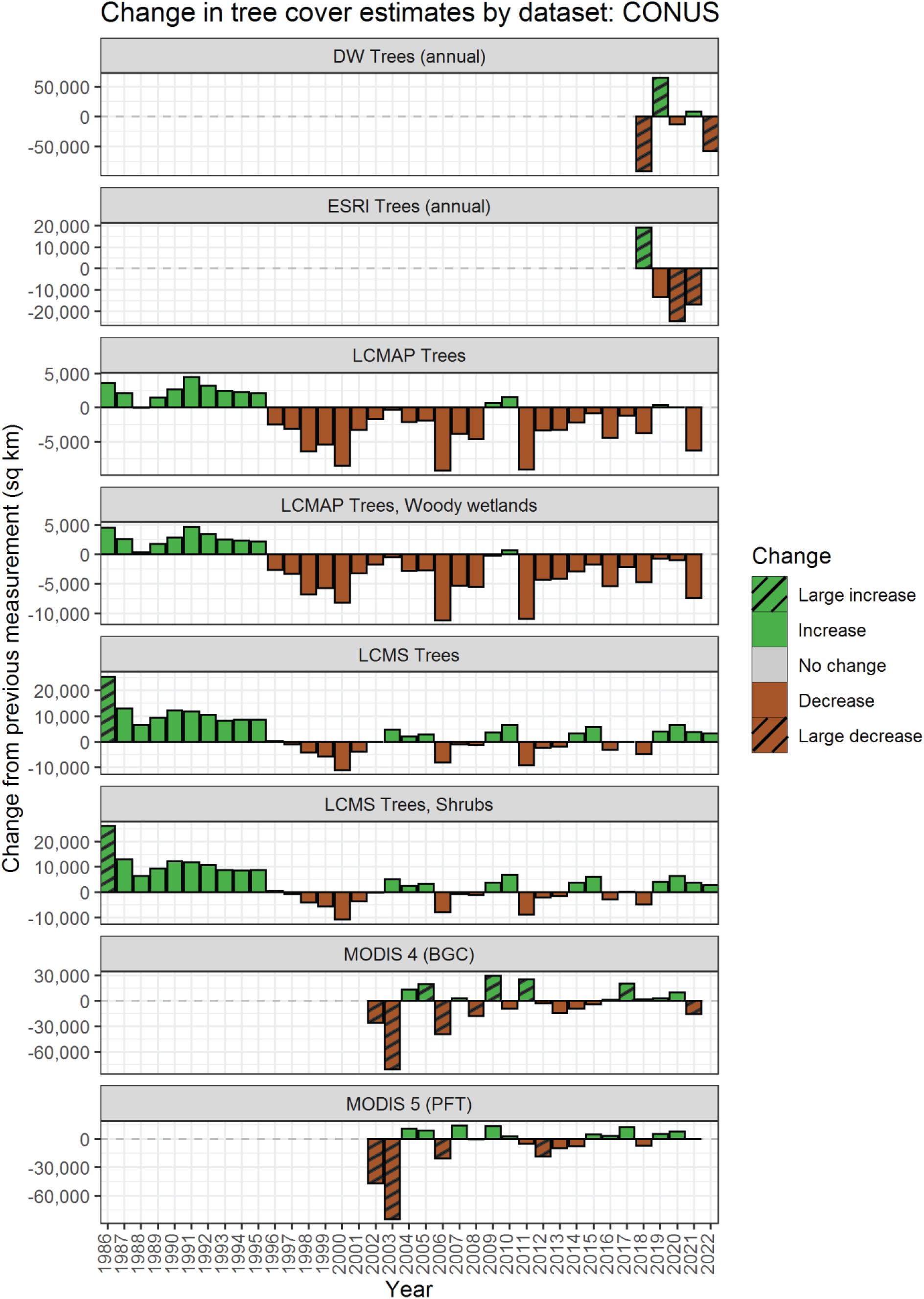
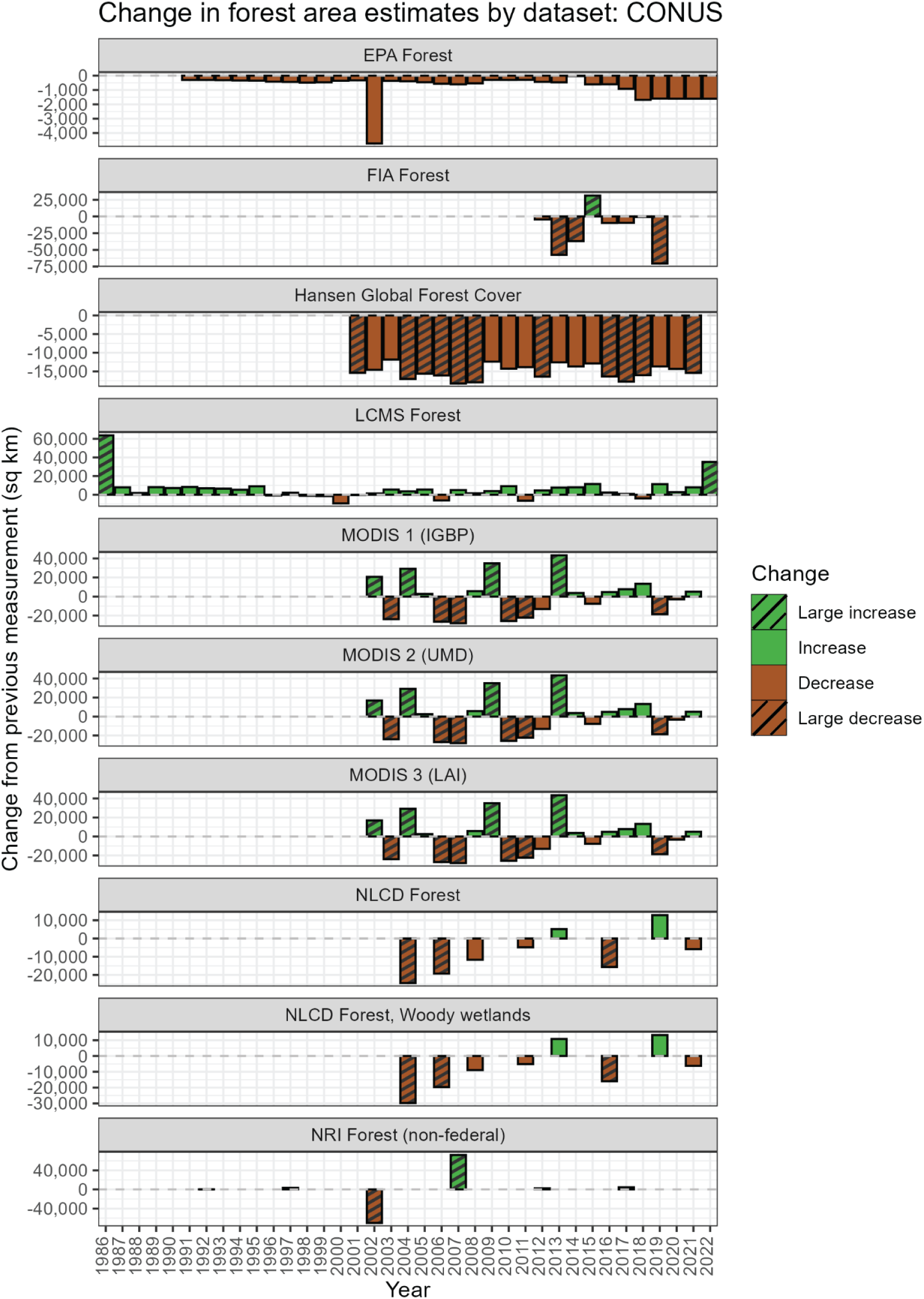
Estimates of 1) tree and 2) forest cover normalized to show change from the previous measurement. “Large” magnitude changes (>15,000 sq km) have stripes to help orient the reader to different y-axis scales. There was no change between 2021-2022 in the ESRI Trees dataset.

Some years show significant discrepancies in direction of change across datasets. For example, FIA Forest shows a loss of ∼57,000 km^2^ of forest from 2012-2013. Over this same year, MODIS 1, MODIS 2, and MODIS 3 reported a gain of ∼43,000 km^2^ (Fig. 2). From 2018 to 2019, FIA Forest reported a loss of ∼70,000 km^2^, while Dynamic World Trees reported a gain of ∼64,000 km^2^ (Fig. 2). Such conflicts between datasets over the same time period reinforce the importance of choosing data with the definitions and methods most appropriate for a given project. There is no single explanation for these discrepancies – these datasets have different definitions (Table 1), spatial resolutions (500 meters versus 10 meters), and methods, as FIA is an inventory-based land-use dataset, MODIS produces annual remotely sensed land-cover classifications using decision trees, and Dynamic World produces near-real-time remotely sensed land-cover classifications using a fully convolutional network. Any number of dataset characteristics can contribute to patterns that differ from patterns in other datasets.

### Mixed trend results at CONUS and state scales

We conducted linear trend analysis from 2000-2019 for datasets with at least four observations within this period (see Methods). Of the 18 datasets included in the trend analysis (see Fig. 4), two have significant increases, ten have significant decreases, and six have no significant change from 2000-2019 at the CONUS scale (Fig. 1; Fig. 5 inset map). The two datasets with significant increasing trends in forest are LCMS Forest (land use) and LCMS Trees, Shrubs, and Barren (land covers). The similarity in estimates and trends among these two datasets is perhaps because the three combined land covers encapsulated in the latter are all part of forest successional patterns following disturbances, such as fire or tree harvesting (i.e., forest use encompasses these land covers). LCMS is the only land cover dataset that shows no significant decreases from 2000-2019 (Fig. 1). In contrast, both NLCD and both LCMAP configurations report significant decreases over this period (Fig. 1). Significant decreases are also found in both FIA datasets and EPA Forest, both inventory-based datasets (Fig. 1).

These disparate trends are reflected in the correlative relationships (Fig. 3). If datasets differed in the magnitude of their area estimate but not the direction of change, we would expect their time series to be significantly and positively correlated. However, only 31% of datasets have significant positive correlations, while 16% have significant negative correlations (Fig. 3). Such mixed correlative relationships may be due to differences in observation methods (e.g., inventory versus remote sensing), definitions (e.g., use versus cover, coverage thresholds, forest type distinction), and spatial resolutions. As an example, the USDA FIA Forest dataset and the US EPA Forest dataset are both positively correlated with each other, and negatively correlated with the USGS LCMS Forest dataset (Fig. 3). Forest estimates in EPA data are primarily based on FIA (U.S. Environmental Protection Agency, 2023), while LCMS does not use FIA data and instead relies on remote sensing images and change detection algorithms (Housman, Heyer, et al., 2023).

**Figure 3.**
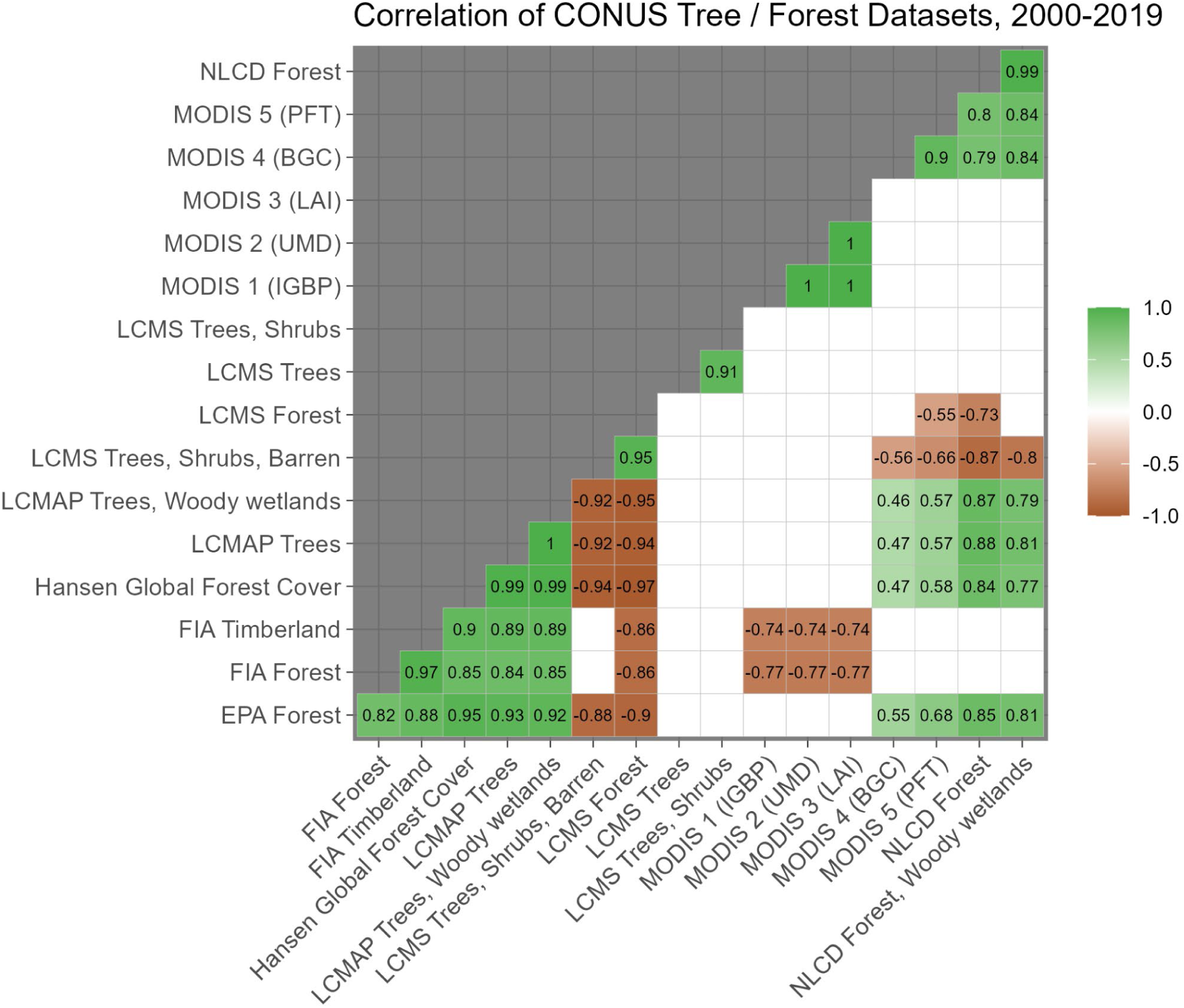
Correlation matrix of forest and tree cover datasets for the conterminous United States from 2000-2019. Only statistically significant correlations are shown. Insignificant correlations are empty white squares. NRI is not shown here because of insufficient data to calculate significance.

At the state level, trend direction and significance vary substantially among datasets and are sensitive to differences in land use versus land cover and definitions of what constitutes a forest (Fig. 4). Most states have more decreasing trends than increasing or insignificant trends in forest or tree cover, and 16 states have majority decreasing trends (Fig. 5). Most agreement among those with decreasing trends occurs in western and northeastern states (Fig. 5). Several states, primarily in the midwest and south, have more increasing trends than decreasing trends, although agreement across all datasets in these states is not as strong as in states with mostly decreasing trends (Fig. 5). Some states show many insignificant trends over time, notably Texas, Oklahoma, Maine, Tennessee, and North Carolina, which show no significant trend for more than half the datasets analyzed (Fig. 5). Arkansas, Delaware, Colorado, Connecticut, West Virginia, and Montana have three or fewer insignificant trends (Fig. 5), although this does not mean that there is agreement in the direction of significant trends (e.g., Arkansas [AR] in Fig. 5).

**Figure 4.**
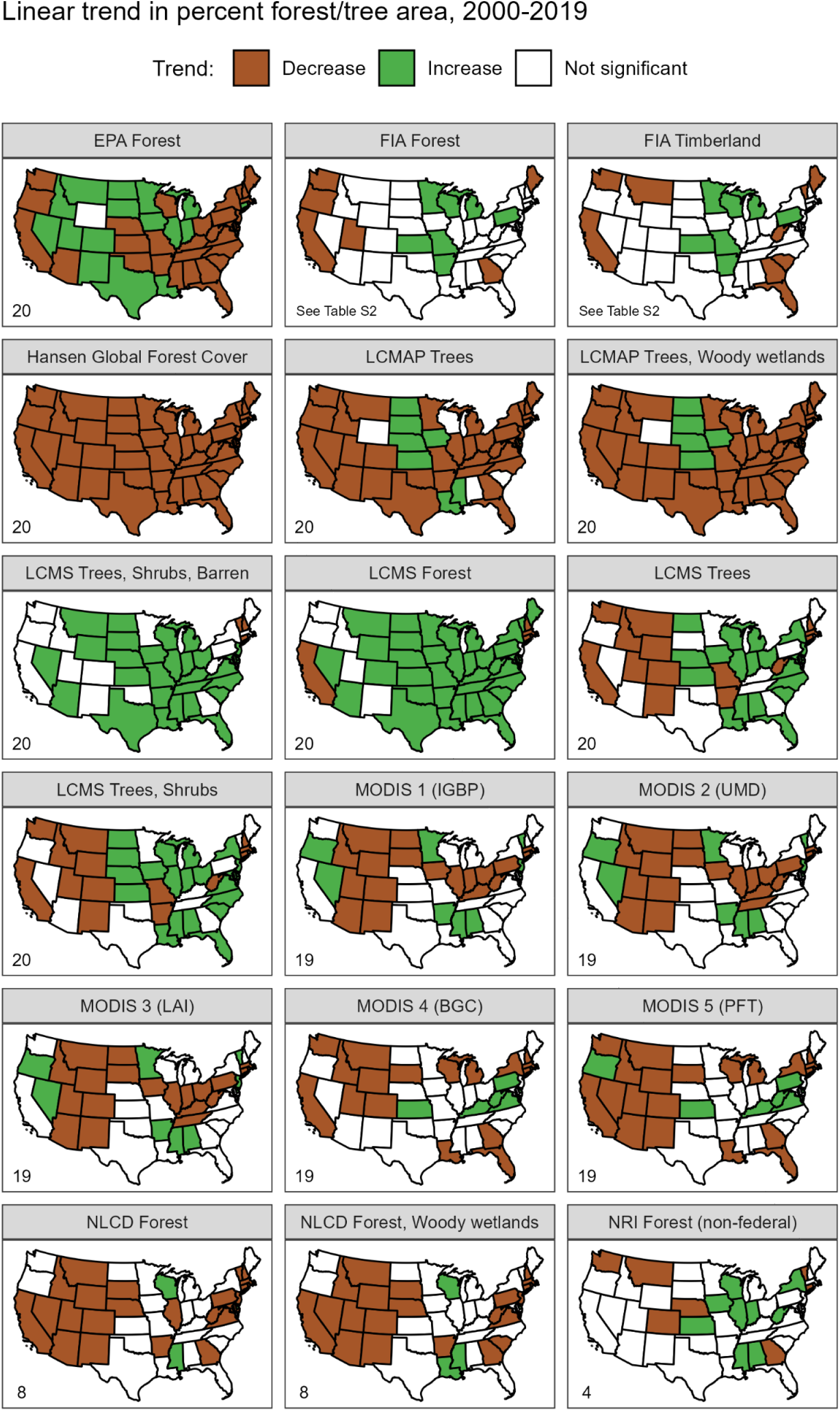
Linear trend results of percent forest or tree area over time (2000-2019) by dataset and state. Numbers in lower left corners indicate number of observations.

**Figure 5.**
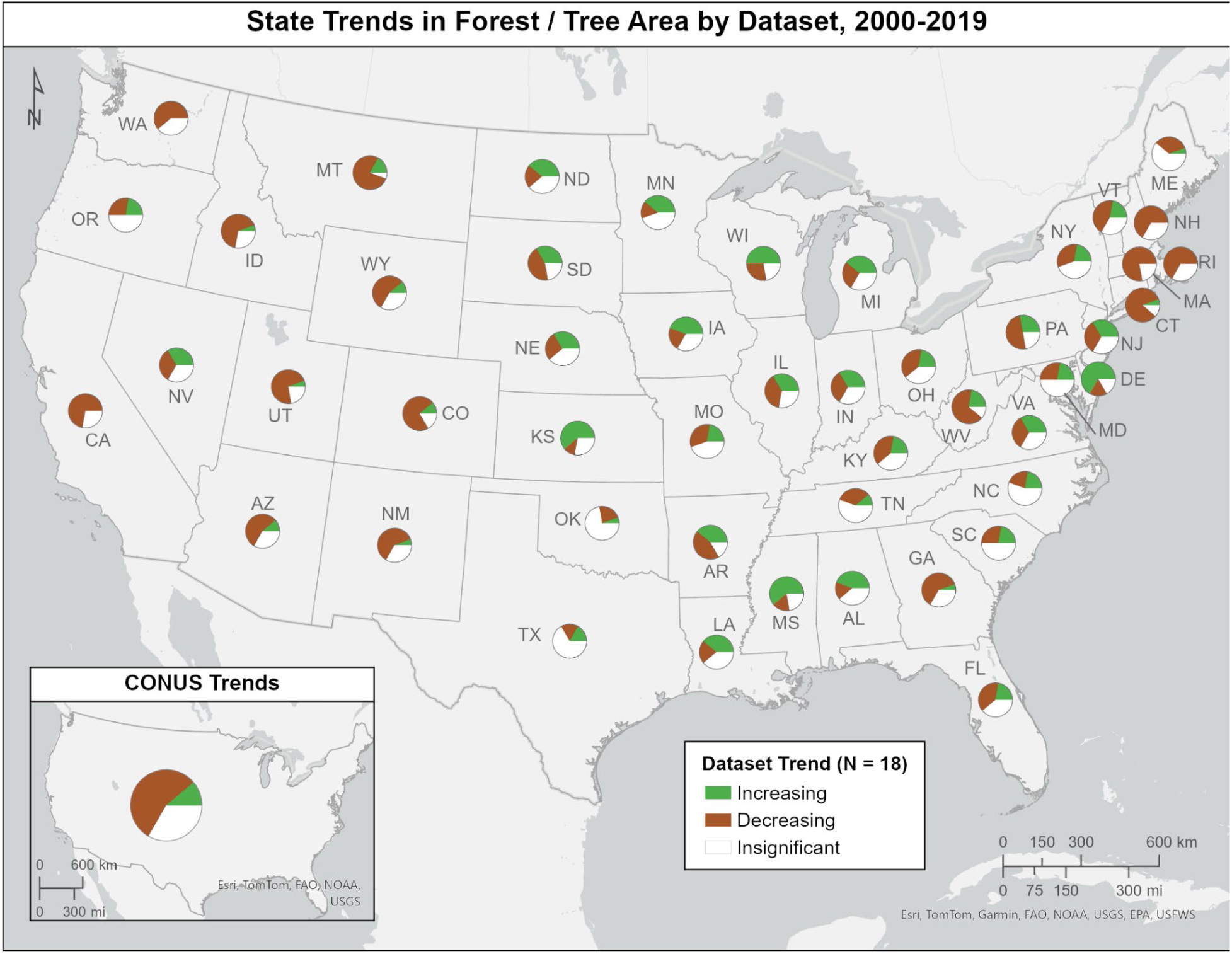
Map summarizing the significant and insignificant trends in forest or tree area from 2000-2019 by state. Pie charts show the number of increasing (green), decreasing (brown), and insignificant (white) trends in each state (main map) and for CONUS (inset) among the 18 datasets analyzed.

## Discussion

Our results show the degree to which forest datasets disagree and reinforce the importance of making informed choices when selecting datasets for analysis. Indeed, someone wishing to understand the status of forests in the U.S. might reasonably conclude that CONUS has lost 81,991 km^2^ of forest over the past 30 years (if they choose LCMAP Trees and Woody Wetlands; Fig. 1). Alternatively, they could just as reasonably conclude that CONUS has gained 93,536 km^2^ (if they choose LCMS Forest; Fig. 1). In the following discussion, we elaborate on the characteristics we highlighted in the sections above and provide some practical guidance for consideration during the dataset selection process.

While the following section provides a deep dive into dataset characteristics and selection, our comparison shopper’s guide (Fig. 6) provides a high level summary that can be used as a quick reference guide. We also developed online tools to facilitate data selection and exploration. The online comparison shopper’s guide (https://ee-forests.projects.earthengine.app/view/forests-shopper) is an interactive tool to filter down dataset options based on desired characteristics, and the forest dataset explorer (https://valeriepasquarella.users.earthengine.app/view/forest-dataset-explorer) provides an interface to explore all of the datasets analyzed here.

**Figure 6.**
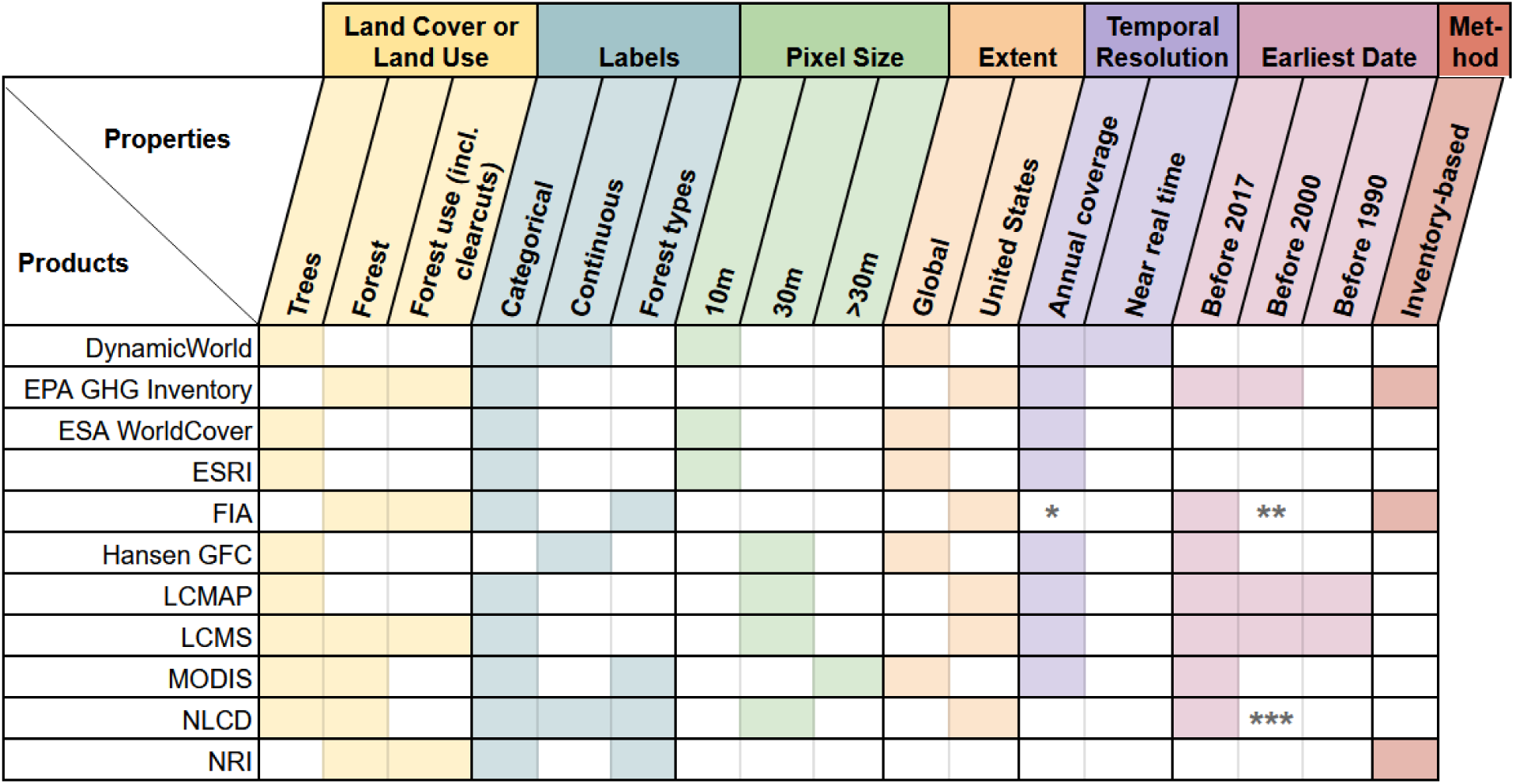
Comparison shopper’s guide summarizes characteristics commonly considered prior to choosing a dataset for an analysis. *FIA plot remeasurement does not occur annually. However, statistical methods can be used to create annualized data such as with the rFIA package. **FIA began prior to 2000 but used a different measurement protocol and may not be comparable to data collected 2000 or later. ***NLCD has a dataset for 1992 but it is not comparable with later years.

### Definitions and characteristics

#### Land use or land cover

As a starting point, the savvy comparison shopper should first ask whether they are seeking to understand forest as a use or as a cover (Table 2, Fig. 6). Whether forests are defined as a land use or land cover has a strong influence on estimates of area and of change, especially in regions where there is extensive and intensive tree harvesting (Coulston et al., 2014; Holmgren, 2015; Woodall et al., 2016). The fundamental difference between forest-use and forest-cover datasets is whether lands absent of trees can be considered forest. Therefore, forest-use datasets may not be appropriate for questions of forest health or degradation (Chazdon et al., 2016), as they overlook fragmentation due to harvesting and do not correspond to the presence or condition of trees on the landscape. Forest-cover datasets also overlook differences in structure and composition in forests that are relevant for biodiversity and conservation (Chazdon et al., 2016). On the other hand, forest-use data collected in the field often include additional attributes that could aid in analysis. In the US, FIA data contains tree presence/absence, species, and biomass information that makes it a commonly used dataset for estimating carbon stocks and is considered useful for applications related to forest health (Tinkham et al., 2018). FIA also distinguishes timberlands from forests (Table 1), allowing for ecological and economic analysis of timberlands. Trend analyses for FIA Forest and FIA Timberland were not identical, and several states showed more declining trends for timberlands than for forest (Fig. 4).

When deciding between land-use and land-cover definitions of forest, a data user can consider whether they are concerned with the landscape as it materially exists or its potential (e.g., the difference between existing forest biomass versus potential to sequester carbon over time). This distinction is especially important in locations where trees are harvested and there may be significant areas without trees that are still in forested use (Fig. 7A). If a user has decided on a land-cover dataset, the next question is what land-cover classes are available and how these relate to the application of the data and the land covers in the study area. This can be particularly important in places with substantial shrublands or woody wetlands, which may be classified as forest or another category (Fig. 7B, 7D). Forests mapped with and without woody wetlands affected trends results in some states (Fig. 4) and has been found to influence assessments of change over time in other studies (Coulston et al., 2014). Whether information of forest types is important can also limit options (Table 1, Table 2, Fig. 6).

**Figure 7.**
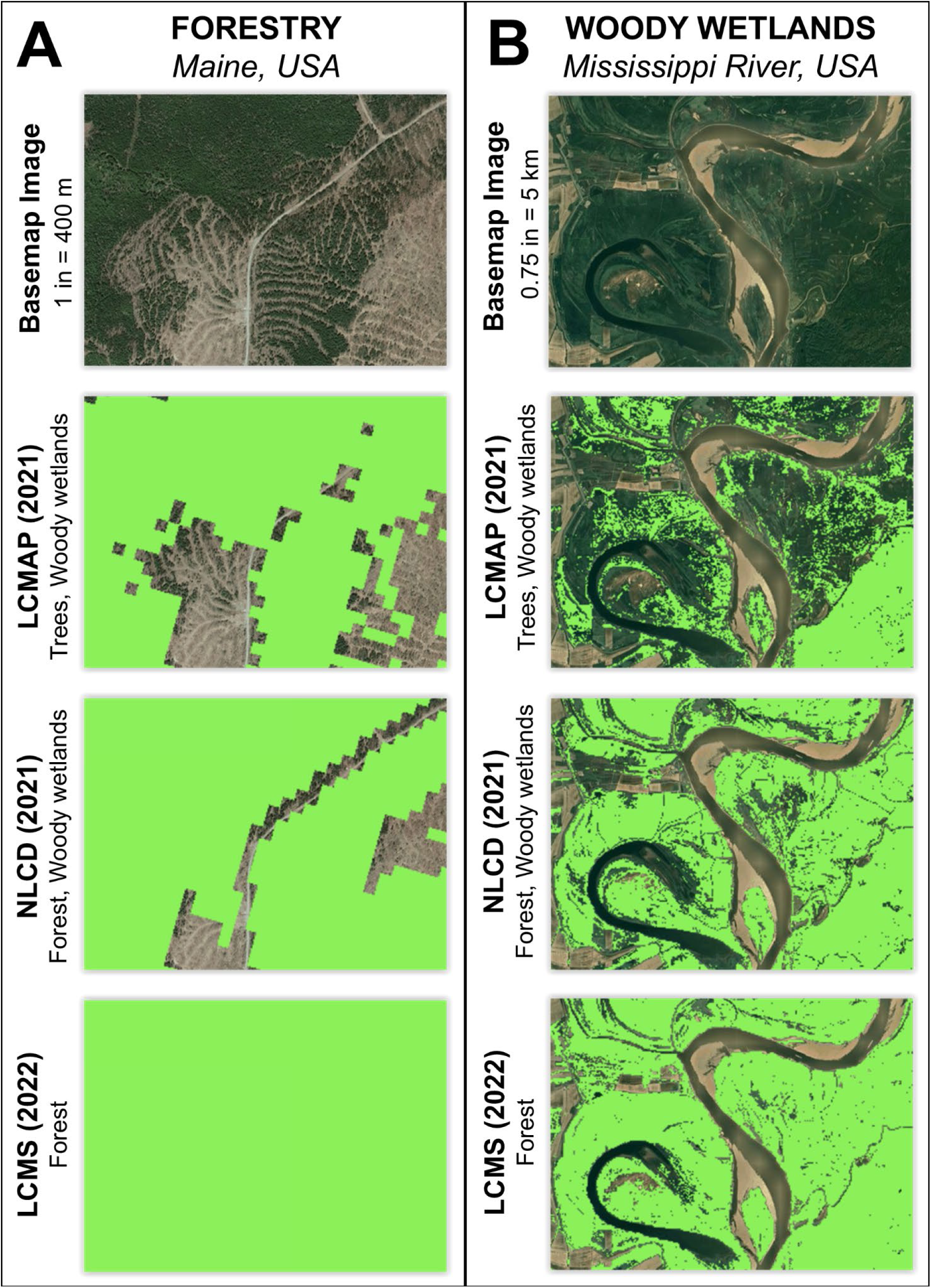
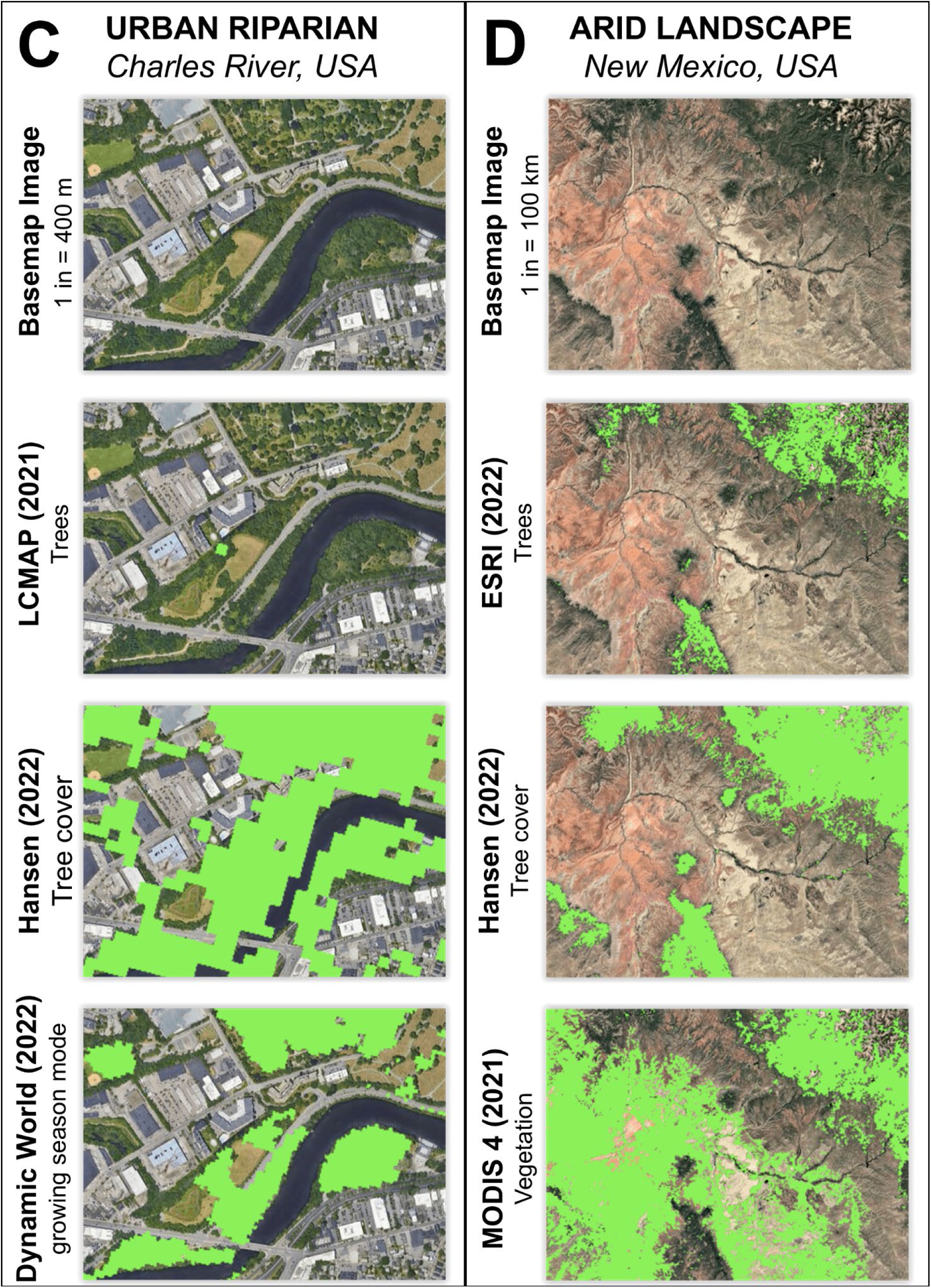
Examples of datasets in different landscapes and scales. **A)** A rural forestry-dominated area, with three examples showing 30 m forest cover (top, middle) and forest use (bottom). **B)** A riparian area with woody wetlands and three examples showing 30 m forest cover (top, middle) and forest use (bottom). **C)** An urban riparian area with small clusters of trees and examples showing two 30 m land cover datasets with diverging maps and a 10 m land cover dataset. **D)** An arid landscape showing land cover datasets with different canopy height requirements (15 m - top, 5 m - middle, 1 m - bottom).

An important part of the definitions of forest-cover and forest-use datasets is the threshold at which something becomes “forest”. Both forest-cover and forest-use datasets employ thresholds that may not capture policy-relevant forest changes (Zalles et al., 2024). For example, if policy dictates that land is forested if there is 10% tree coverage (the most common threshold used in the datasets analyzed here, see Table 1) over a certain size area (whether a 30 m pixel or a 0.4 ha plot), any gain greater than 10% within that area is not documented in a categorical map. Nor would a loss from 90% tree cover to 20% tree cover be documented, although such a change would have implications for policies related to forest loss or conservation. In contrast to categorical maps where “forest” or “tree” is defined and mapped for you, continuous datasets (Table 2, Fig. 6) provide more flexibility because the user can choose their own threshold by which to determine what counts as “tree” or “forest”.

Both forest-cover and forest-use datasets that aggregate monoculture plantations and native forests will obscure the diminished benefits of plantations (Van Holt et al., 2016), and may overestimate wildlife habitat, biodiversity, value for Indigenous peoples, or other dimensions of forests that depend on species composition and structure (Chazdon et al., 2016). While forest-cover and forest-use datasets may not contain information on species, structure, biodiversity, and health that are important for many conservation-related questions and related policy, other datasets can be used to fill in these gaps. NLCD land cover can be combined with the NLCD percent canopy cover which helps estimate canopy density and structure (Sohl et al., 2025). NDVI is a remote-sensing based index often used to approximate vegetation health (e.g., Barka et al., 2019; Z. Wang et al., 2021). Imputed maps of forest composition and structure can also be used to represent these variables integrated with or instead of cover maps (Duveneck et al., 2015; Ohmann & Gregory, 2002). Management (i.e., use) is difficult to determine through remote sensing alone, but there have been efforts to map forest management in recognition of its substantial impact on forests (Lesiv et al., 2022).

#### Temporal resolution and depth

Another key consideration for a forest data user is temporal resolution (frequency of observations) and depth (duration of the time series). The time period of analysis also constrains data options. Only LCMAP, LCMS, and NRI provide data prior to 1990 (Table 2, Fig. 6, Fig. S1). The temporal depth of these datasets has led to their use in broad scale studies quantifying multidecadal land cover trends and variation in land cover changes over time (e.g., Auch et al., 2022; Bigelow et al., 2022; Dwomoh & Auch, 2024). EPA Forest began in 1990 and provides a forest land-use time series of nearly 25 years (Fig. 1). Analyses focused on the most recent two decades have more options, as many widely used datasets including FIA, NLCD, Hansen, and MODIS became available in the 2000s (Table 2, Fig. 6). These datasets also have multiple decades of data and are similarly used for trend analysis of land-cover change and of vegetation specifically (e.g., Hansen et al., 2013; Homer et al., 2020; Nelson et al., 2020; Zhu et al., 2021). Despite using input sensors with different temporal revisits, most datasets summarize forest conditions over an entire year. Applications in which near-real-time data is valuable may be best-suited by datasets like Dynamic World, which is generated for individual Sentinel-2 images, while applications that require calibration with field samples may require use of inventory datasets, despite longer revisit times (Table 2, Fig. 6).

There is often a tradeoff between temporal resolution and classification detail. For example, NLCD has four development classes and three forest classes but has historically only been available every 2-3 years. NRI, which has the most detailed classification scheme of any dataset analyzed here, is only available every five years. In contrast, LCMAP and LCMS have relatively simple classification schemes (e.g., one development and one tree class) but provide data every year over almost 30 years. The particulars of the analysis can help inform whether it is better to compromise on temporal precision or classification detail. This long standing tradeoff is also being mitigated by the maturation of technology and data production. For example, in late 2024 the US Geological Survey merged NLCD and LCMAP to create a new dataset that combines the thematic detail of NLCD with the annual temporal resolution of LCMAP (Sohl et al., 2025).

#### Spatial resolution and scale

The spatial resolution of a dataset has a significant influence on estimates of spatially-dependent characteristics such as forest fragmentation (Morreale et al., 2024; Wickham & Riitters, 2019) and connectivity (Hernando et al., 2017). For forest-cover datasets where forest is classified based on percent of a pixel covered by tree canopy, spatial resolution is a critical consideration and should be considered in the context of study area location and scale of analysis, as a spatial resolution too large can obscure real world patterns (Fig. 7A) and a resolution too small can be computationally infeasible. Wang and Mountrakis (2023) assessed the accuracy of many datasets analyzed here and concluded that LCMAP and NLCD performed the best at CONUS scale, but that there were regional variations in accuracy for every dataset. Datasets that perform well in a certain location at a large scale may not be the best choice in a subregion or at a smaller scale. For example, LCMAP appears unable to detect trees along an urban river in Massachusetts (Fig. 7C).

In this analysis, most remote sensing datasets come from the Landsat family of satellites and are at 30 m resolution, which is considered moderate resolution (Table 2, Fig. 6). While spatial resolution is an important consideration, our results show that spatial resolution alone is not enough to explain variation in forest area or trend estimates (Fig. 1). For example, among 500 m MODIS datasets, MODIS 1-3 estimate the lowest forest area of datasets analyzed while MODIS 4 estimates the highest area (Fig. 1). Similarly, among 30 m datasets, Hansen estimates less than half the forest area that LCMS estimates in 2020 (Fig. 1). While choosing an appropriate spatial resolution is important, even among datasets that share spatial resolutions, differences in definitions, classification, and change detection algorithms can lead to disparate results (Chen et al., 2020).

Forest-use datasets based on inventory assessments are typically at a coarser spatial resolution than forest datasets created with remote sensing. Publicly available FIA data is “fuzzed” and the precise location of individual field plots is not available without special permission, which complicates a user’s ability to integrate with remote sensing data, especially at finer spatial resolutions where fuzzing could impact analyses (Tinkham et al., 2018). Analyzing FIA data also has a significant learning curve (Tinkham et al., 2018). NRI and EPA, the other inventory-based forest use datasets, are not available below the county or state level, respectively, which limits their utility to large scale analyses. LCMS, the only remote sensing forest-use dataset in this analysis, may be an alternative for questions of forest land use at sub-county scales if using FIA data is not feasible.

#### Area and height requirements

Thresholds for the area of forest and the height of vegetation are common components of forest classifications (Table 1). Awareness of how such requirements include and exclude certain segments of forests is helpful when using forest datasets. Area requirements are a feature of forest-use datasets created through forest inventories (Chazdon et al., 2016). All three inventory-based datasets analyzed here – EPA, FIA, and NRI – require a minimum area of 0.4 ha and width of 100 or 120 ft to be considered “forest” (Table 1). Such requirements may exclude trees in urbanized landscapes, narrow strips of trees in riparian or agricultural areas, and remnant trees (Chazdon et al., 2016), all of which are important forest areas for conserving biodiversity (Arroyo-Rodríguez et al., 2020).

Height requirements are present in both inventory-based and remote sensing-based forest datasets (Table 1). Such requirements may exclude certain tree species or early successional forest. ESRI has the tallest canopy height threshold at 15 m, which includes only mature canopy trees in the US, while MODIS-derived classifications have the lowest canopy height threshold at 1-2 m and may include shrubs (Table 1). Differences in canopy height requirements can produce substantially different “forest” maps in locations where vegetation height is shorter or more variable, such as in regenerating forest, chaparral and/or arid landscapes (e.g., Fig. 7D). Most datasets analyzed have a canopy height requirement of 4-5 m (Table 1), which in the US includes young canopy trees and smaller tree species, but not very early successional forest. Such height requirements may help distinguish forest from shrub in certain landscapes but may also lead to underestimating reforesting or afforested areas (Chazdon et al., 2016).

### Change over time estimations and algorithms

How forest datasets estimate change in forest area over time is related to their definitions, scales, and change detection methods. The role of time is treated differently when estimating forest use than for forest cover, as forest use datasets are less affected by short term land cover changes and aim to represent management intent over successional time frames (Coulston et al., 2014; Winkler et al., 2021). In this analysis, forest land-use and forest land-cover datasets have similar proportions of datasets estimating no significant change or a significant decrease (2 of 4 for land-use datasets and 6 of 13 for land-cover datasets; Fig. 1). However, only one of thirteen forest land-cover datasets suggests a significant increase compared to one of four land-use datasets (Fig. 1). Additionally, the EPA Forest land-use dataset, while showing a statistically significant decrease in forest area, has an extremely low magnitude of change (Fig. 2) and appears “stable” in the context of other datasets (Fig. 1). In addition to differences in definitions of forest as use versus cover, the scale of analysis influences detection of change over time, which presents a challenge given that analyses are typically conducted at a single scale and change occurs over multiple spatial and temporal scales (Coulston et al., 2014).

With multiple decades of remote sensing data available to provide reference data and the development of robust algorithms, many datasets consider change over time directly in the data creation process in order to make data more accurate (e.g., less susceptible to spurious changes) and more comparable to itself over time. For example, NLCD produces its datasets in suites of harmonized datasets that are directly comparable over time (Jin et al., 2019). LCMAP uses the Continuous Change Detection and Classification (CCDC) algorithm to produce its annual land cover maps (US Geological Survey, 2022), while LCMS uses CCDC and the LandTrendr algorithm to do so (Housman, Heyer, et al., 2023). Pasquarella et al. (2022) describe the key characteristics of and differences between these algorithms that can help users of these datasets gain a deeper understanding. More broadly, when estimating change in forest area over time from classified maps, users should seek out information on the comparability of the data over time and any recommended steps to account for bias or inaccuracy in area estimations. While in some cases dates may be directly comparable, in other instances the metadata may provide bias correction constants that can be applied to account for inaccuracies in the map classifications (Olofsson et al., 2016).

Standard practice in change detection is to use probability-based samples of reference data rather than directly comparing maps (Olofsson et al., 2014). Such methods require significantly more time, and therefore money, to complete and are subject to errors in interpreter bias and discrepancies in spatial and temporal resolution of the map and reference data (Olofsson et al., 2014; Zalles et al., 2024). With the increasing abundance of moderate and high resolution satellite imagery, such methods can also lead to a loss of precision in estimating forest changes compared to using the satellite imagery directly (Zalles et al., 2024). Furthermore, working directly with field inventories, such as FIA, comes with its own challenges in terms of noise and quality of resulting area estimates. These challenges lead many users of forest data, apart from professional researchers, to simply use the datasets as they are. This reality, plus the fact that many forest datasets are interdependent, lead us to compare the datasets directly in “map space” – the actual outcome that is produced as a result of each dataset’s many characteristics.

The interdependence of forest datasets often relates to training data or creating composite data from the best available existing source. For example, the Dynamic World and ESRI datasets share the same training data but use different neural networks for classification (Venter et al., 2022). Both LCMAP and LCMS use NLCD in creating their training data (Housman, Heyer, et al., 2023; Zhou et al., 2020). EPA Forest is a compilation of multiple datasets analyzed here including FIA, NLCD, and NRI (Sohl et al., 2025). All datasets come with a technical report, and we have provided links to those for the datasets used in this analysis (Table 1). While content is often largely technical, we encourage data users to read what they can and ask questions about dependencies and other major methodological characteristics prior to beginning analysis.

### Decision support

In this paper, we have directly compared the estimates (Figs. 1, 2, S1, S2), trends (Fig. 4, 5), and relationships (Fig. 3) within and among numerous forest datasets. We provide context that can help explain the observed differences and inform suitability for particular uses, including dataset definitions (Table 1) and characteristics (Table 2, Fig. 6). To further support data users in the data selection process, we provide online tools that allow users to interactively explore forest maps and view extent estimates by state.

Our tools (https://valeriepasquarella.users.earthengine.app/view/forest-dataset-explorer; https://ee-forests.projects.earthengine.app/view/forests-shopper) allows users to explore and compare all datasets analyzed in this paper. We provide tools with and without filters so the user can narrow down the options of datasets based on their specific application, including: study region, time period, spatial resolution and other characteristics included in Figure 6. Together, these tools provide the capability to explore as many or as few datasets as the user desires. We also make our Earth Engine and other processing code available for extension and application to other regions and/or domains (https://github.com/valpasq/ee-forests/tree/main; https://github.com/hf-thompson-lab/ee-forests-shopper).

## Conclusions

Our results show how important dataset selection can be, and the many ways forest datasets diverge in their estimates. For many projects, there is no perfect dataset, but considering dataset definitions (Table 1) and characteristics (Table 2, Fig. 6) along with the purpose and needs of the analysis can help identify a defensible choice, and interactive tools can be used to see and compare different options in a study area before making a decision. We hope that these tools and the ideas presented in this paper help forest data users become more critical and discerning about the forest data they use for particular projects and questions.

## Supplementary Materials

**Figure S1.**
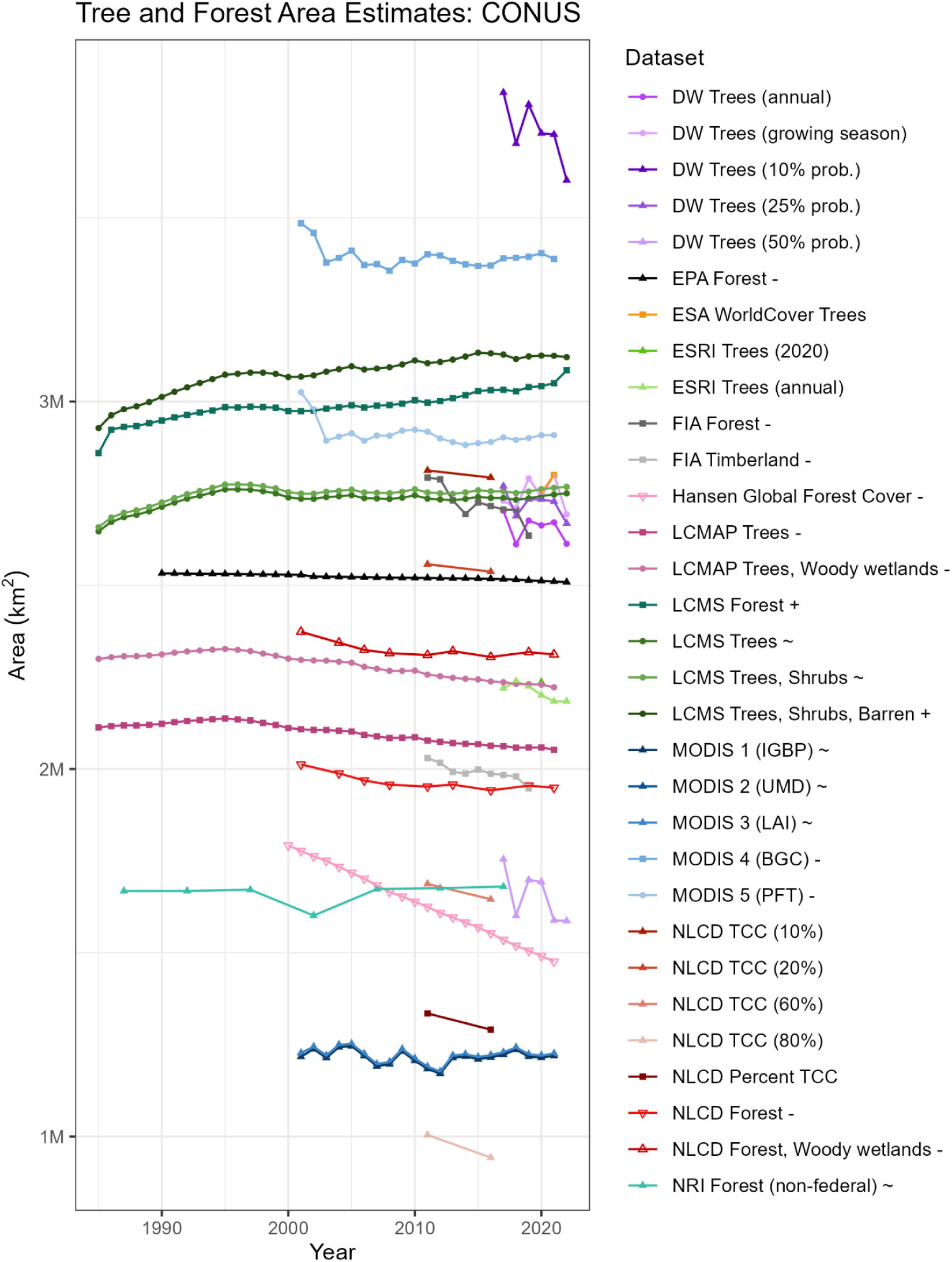
Forest and tree cover area for CONUS, including NRI which excludes federally owned forest. Symbols next to dataset names indicate that the dataset was analyzed for trends and had an increasing (+), decreasing (-), or insignificant (∼) trend from 2000-2019.

**Figure S2.**
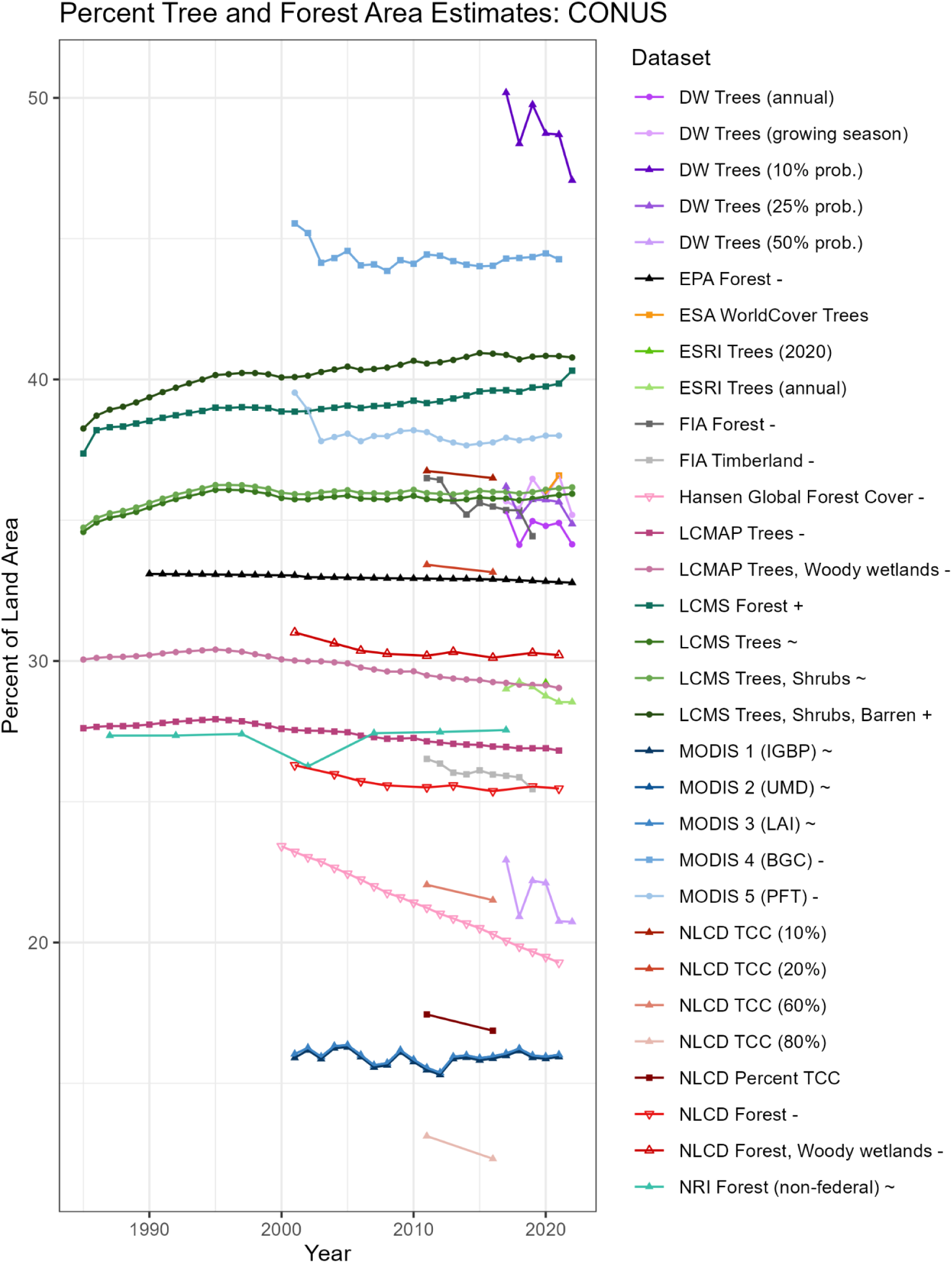
Percent forest and tree cover area for CONUS, including NRI which excludes federally owned forest. Symbols next to dataset names indicate that the dataset was analyzed for trends and had an increasing (+), decreasing (-), or insignificant (∼) trend from 2000-2019.

**Table S1.**
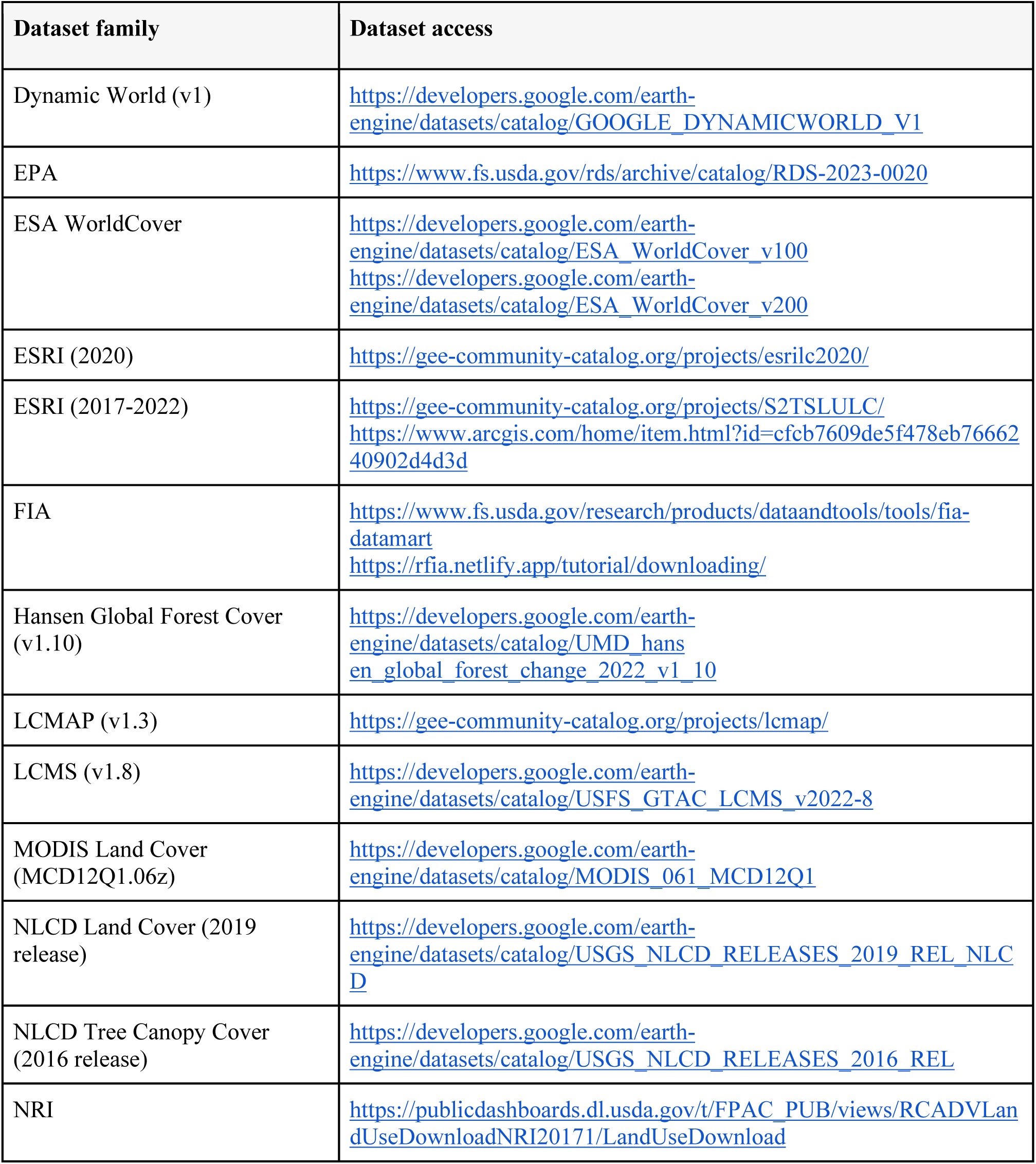
Links to access the datasets analyzed.

**Table S2.** Number of observations used for trend analysis (2000-2019) of FIA data.

| State name (abbreviation) | Number of observations |
| --- | --- |
| Contiguous United States (CONUS) | 9 |
| Alabama (AL) | 19 |
| Arkansas (AR) | 19 |
| Arizona (AZ) | 19 |
| California (CA) | 19 |
| Colorado (CO) | 18 |
| Connecticut (CT) | 17 |
| Delaware (DE) | 16 |
| Florida (FL) | 16 |
| Georgia (GA) | 20 |
| Iowa (IA) | 20 |
| Idaho (ID) | 16 |
| Illinois (IL) | 19 |
| Indiana (IN) | 20 |
| Kansas (KS) | 19 |
| Kentucky (KY) | 20 |
| Louisiana (LA) | 17 |
| Massachusetts (MA) | 17 |
| Maryland (MD) | 16 |
| Maine (ME) | 20 |
| Michigan (MI) | 20 |
| Minnesota (MN) | 20 |
| Missouri (MO) | 20 |
| Mississippi (MS) | 12 |
| Montana (MT) | 17 |
| North Carolina (NC) | 17 |
| North Dakota (ND) | 19 |
| Nebraska (NE) | 19 |
| New Hampshire (NH) | 18 |
| New Jersey (NJ) | 16 |
| New Mexico (NM) | 15 |
| Nevada (NV) | 16 |
| New York (NY) | 18 |
| Ohio (OH) | 19 |
| Oklahoma (OK) | 12 |
| Oregon (OR) | 19 |
| Pennsylvania (PA) | 20 |
| Rhode Island (RI) | 17 |
| South Carolina (SC) | 20 |
| South Dakota (SD) | 19 |
| Tennessee (TN) | 20 |
| Texas (TX) | 18 |
| Utah (UT) | 20 |
| Virginia (VA) | 19 |
| Vermont (VT) | 17 |
| Washington (WA) | 18 |
| Wisconsin (WI) | 20 |
| West Virginia (WV) | 16 |
| Wyoming (WY) | 9 |

## Notes

https://github.com/hf-thompson-lab/forest-comparison

https://github.com/valpasq/ee-forests/tree/main

https://ee-forests.projects.earthengine.app/view/forests-shopper

https://valeriepasquarella.users.earthengine.app/view/forest-dataset-explorer

https://github.com/hf-thompson-lab/ee-forests-shopper

### Competing Interest Statement

The authors have declared no competing interest.

### Funding

- National Science Foundation: DEB-LTER-280 18-32210

